# Cell cycle-dependent transitions in PCP protein mobility erase and restore planar cell polarity in mitosis

**DOI:** 10.64898/2026.01.11.698900

**Authors:** D. Ouyang, L.P. Basta, P. Sil, B.W. Joyce, E. Posfai, D. Devenport

**Affiliations:** Department of Molecular Biology, Princeton University, Princeton, NJ 08544

**Keywords:** planar cell polarity, mitosis, cell cycle, Plk1, Fz6, Vangl2, Celsr1

## Abstract

Central to our understanding of how cells establish polarity is deciphering how polarity proteins are transported to and retained at asymmetric positions. Here, in the context of planar cell polarity (PCP) remodeling in mitosis where asymmetry is transiently lost and restored, we show that PCP proteins transition between mobile and stable states in coordination with cell cycle progression. Live imaging and FRAP analyses reveal that, in addition to becoming internalized into endocytic vesicles, membrane-associated PCP proteins dramatically increase their lateral mobility upon mitotic entry in a Plk1-dependent manner. Unexpectedly, asymmetry is not restored in cytokinesis via polarized delivery of PCP-protein containing vesicles. Rather, PCP proteins continue to diffuse laterally during cytokinesis where they are captured and stabilized non-cell autonomously through adhesive interactions with neighbors. We propose that a transient, laterally diffusive state in mitosis allows for flexibility and robustness in planar polarity as dividing cells change shape and rearrange.

## INTRODUCTION

Cell polarity is a universal feature of cells in which organelles, cytoskeletal structures and membrane components are distributed asymmetrically. Establishment and maintenance of polarity are essential for cellular functions such as directed migration, asymmetric cell division, and tissue morphogenesis^1–4^. Several polarity pathways have been identified and although they consist of distinct protein complexes, they share a universal feature in which membrane-associated polarity proteins are asymmetrically localized to opposing regions of the plasma membrane where they act on downstream factors to direct polarized cell behaviors^5–8^. Deciphering how polarity proteins are transported to and retained at their asymmetric positions is central to our understanding of how cells establish and regulate a polarized state.

As polarity proteins typically begin uniformly distributed, polarization requires transport of opposing polarity proteins to attain asymmetric localizations. Movement can occur through passive diffusion or active transport mechanisms, but once established, movement must be restricted so that asymmetry is maintained^9^. Polarity proteins can be trapped and stabilized by a variety of mechanisms, for example by forming large, higher order assemblies through clustering, becoming anchored to cytoskeletal structures or, in the case of transmembrane polarity proteins, by latching onto an extracellular tether^9^. Understanding polarization therefore requires defining the mechanisms that control how polarity proteins transition between mobile and stable states. In some epithelia, cell polarity is temporarily lost and regained when cells divide^10^, offering an opportunity to define the mechanisms by which polarity proteins transition between transport and retention over relatively short time periods.

Planar cell polarity (PCP) refers to cell polarization along a tissue plane and is an excellent polarity system to investigate dynamic shifts in polarity states. In epithelia, planar polarity governs the polarized alignment of cellular protrusions such as insect bristles, stereocilia bundles, motile cilia and mammalian body hairs^4, 11, 12^. It is controlled by a core complex of integral-membrane and associated cytoplasmic proteins that asymmetrically localize at cell junctions. The complex consists of seven-pass transmembrane protein Frizzled (Fz) and its cytoplasmic partner Dishevelled (Dsh; Dvl in vertebrates) on one side of the junction intercellularly associated with tetraspanin Van Gogh (Vang; Vangl in vertebrates) and its partner Prickle (Pk) on the opposing cell interface^13–16^. Both complexes interact with the atypical cadherin Flamingo/Celsr1 (Fmi; Celsr in vertebrates), which engages in homotypic binding and couples opposing PCP complexes between cells (Fig. 1A)^17–21^.

**Figure 1.**
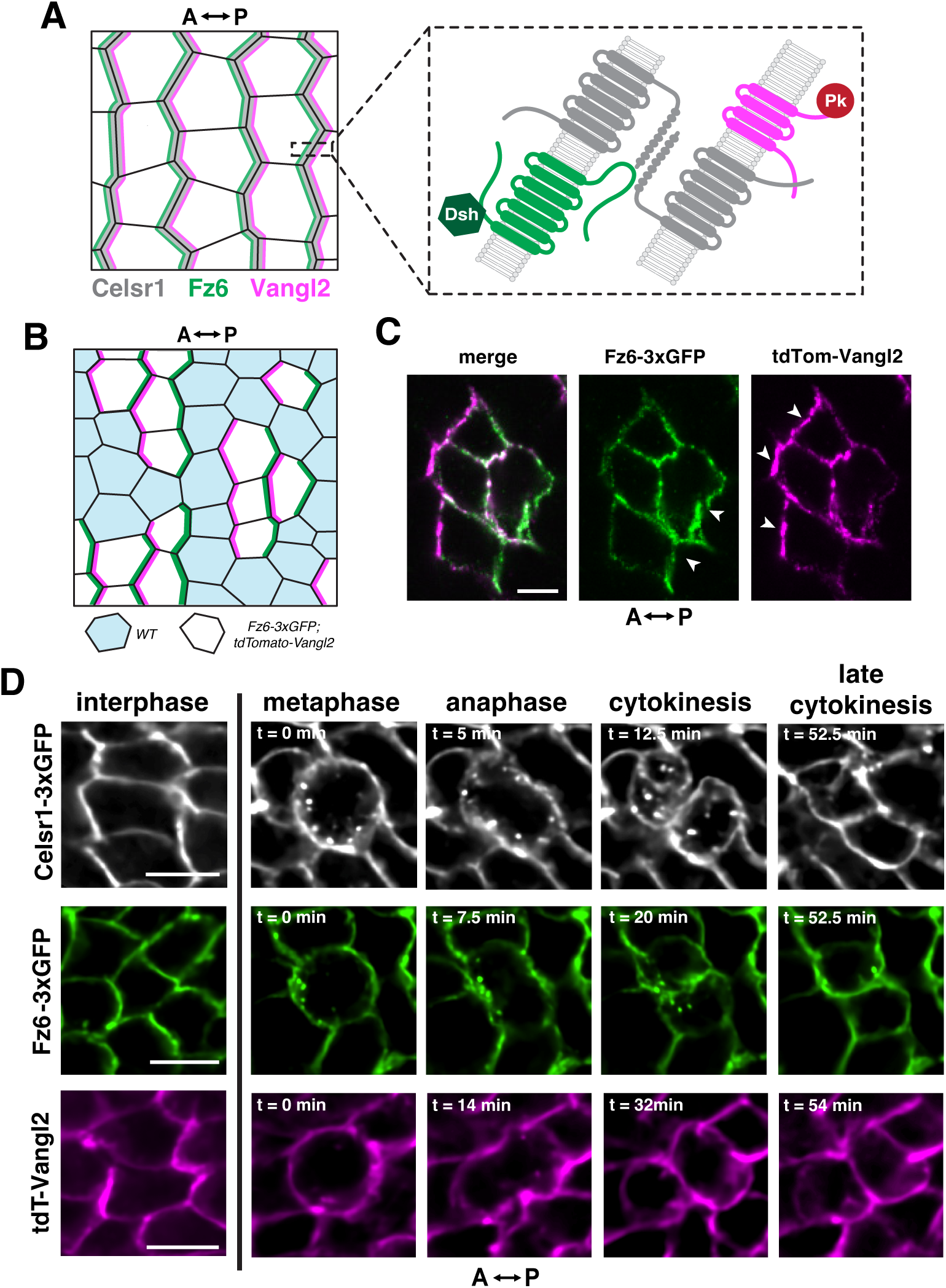
Endogenously-tagged PCP reporter mouse models allow for visualization of PCP proteins in dividing basal epidermal cells. **(A)** Schematic of the planar cell polarity complex in epidermal basal cells. Fz6 (green) and its cytoplasmic partner Dsh (dark green) localize to the posterior cell border opposite of Vangl2 (magenta) and its cytoplasmic partner Pk (red) at the anterior cell border. The atypical cadherin Celsr1 (gray) occupies both borders. **(B)** Schematic of a chimeric epidermis comprised of a mix of dual labeled *Fz6-3xGFP;tdT-Vangl2* cells and *WT* cells. **(C)** Representative planar view of a cluster of dual PCP reporter cells in interphase surrounded by *WT* cells from *E*15.5 *WT : Fz6-3xGFP;tdT-Vangl2* chimeric skin. Arrowheads point to the asymmetric localization of Fz6-3xGFP (green) at the posterior and tdT-Vangl2 (magenta) at the anterior. Scale bar, 5 µm. **(D)** Still frames from live imaging of basal epidermal cells from *E15.5 Celsr1-3xGFP* (top panels) or *Fz6-3xGFP;tdT-Vangl2* (middle and bottom panels) skin explants. Distribution of Celsr1-3xGFP (grayscale), Fz6-3xGFP (green) and tdT-Vangl2 (magenta) at interphase and from metaphase through late cytokinesis. Scale bars, 10 µm.

In proliferative progenitor cells of the murine epidermis, PCP proteins are dynamically remodeled when cells divide such that their asymmetry is transiently erased at mitotic entry and regained in the following interphase^22^. Upon mitotic entry, transmembrane PCP proteins are internalized from the cell surface via endocytosis and held within endosomes until mitotic exit. In cytokinesis, PCP components are returned to the membrane and polarity is eventually restored^22^. This cycle of polarity loss and restoration thus involves destabilization and movement of PCP proteins away from their polarized positions, followed by movement and stabilization to asymmetric positions in daughter cells. We previously showed how cell cycle kinases control the endocytosis and removal of planar polarity proteins upon mitotic entry, but how polarity is restored following cytokinesis remains entirely unknown^23^.

To decipher the mechanisms by which PCP asymmetry is lost and regained, we performed live imaging and FRAP analysis of endogenously-tagged PCP reporters in mouse epidermis to monitor Celsr1, Fz6 and Vangl2 as they are dynamically redistributed in mitosis. We show that mitotic internalization of PCP proteins coincides with a dramatic increase in the lateral mobility of PCP proteins that remain at the plasma membrane, and that this mobility is promoted by the cell cycle kinase Plk1. Consistent with prior data in fixed epidermal tissues, we find that endocytic vesicles containing each of the transmembrane PCP components enter the cell unequally from opposite sides but are partitioned evenly in daughters. Surprisingly, PCP-protein containing vesicles are not redelivered to the membrane in a polarized manner that would restore their asymmetric distributions. Rather, their localization must be further refined following cytokinesis. Using FRAP, we find that PCP proteins continue to be highly mobile within the plasma membrane during cytokinesis and stabilize into regions of high protein density and low protein mobility during interphase. Further, using mosaic approaches with chimeric mouse embryos, we show that capture of PCP proteins by adhesive interactions with neighbors stabilizes them non-cell autonomously. These data demonstrate that PCP proteins transition between mobile and stable states in coordination with cell cycle progression. We propose that a transient, laterally diffusive state in mitosis allows for flexibility and robustness in polarity as dividing cells change shape and rearrange.

## RESULTS

### Endogenously-tagged PCP reporter mouse models allow for visualization of PCP proteins, live, in dividing basal epidermal cells

To visualize how PCP asymmetry is dynamically reorganized across the cell cycle, we took advantage of three mouse strains in which the PCP genes Fz6, and Vangl2 and Celsr1 are fused to fluorescent reporters at their endogenous loci: Celsr1-3xGFP, Fz6-3xGFP, and tdTomato-Vangl2 (tdT-Vangl2)^24^. In interphase, all three PCP proteins localize asymmetrically in basal cells of the skin epidermis, where they enrich at anterior-posterior (AP) junctions, as previously shown^24, 25^. To resolve Fz6 and Vangl2 localization to either side of the junction, we generated mouse chimeras consisting of a mix of wild type, unlabeled and Fz6-3xGFP; tdT-Vangl2 dual reporter cells (Fig. 1B). As expected, and consistent with previous data using mosaic overexpression and super-resolution imaging, Fz6-3xGFP enriches strongly on the posterior side of the cell, while tdT-Vangl2 enriches on the anterior (Fig. 1C)^24^.

Next, we performed live imaging of E15.5 skin explants from Celsr1-3xGFP or Fz6-3xGFP; tdTomato-Vangl2 embryos and followed basal progenitors as they divide (Fig. 1D, Movie S1, Fig. S1). As observed previously by immunofluorescence in fixed tissue samples, endosomes containing Celsr1-3xGFP, Fz6-3xGFP and, to a lesser extent, tdT-Vangl2 were internalized from the cell surface at the onset of mitosis^22, 26, 27^. Additionally, we observed that a significant proportion of Celsr1-3xGFP, Fz6-3xGFP and especially tdTomato-Vangl2 remained at the plasma membrane of dividing cells, which was more evident in live than in fixed epidermal samples^22, 24, 26, 27^.

### Membrane-localized PCP proteins are mobilized during mitosis

In the *Drosophila* wing, asymmetrically localized PCP proteins are highly stable within the membrane, and this immobile state is thought to be key for maintaining asymmetry^28, 29^. Considering that in dividing cells a proportion of PCP proteins is internalized from the cell surface, we hypothesized that such a reduction of PCP-mediated cis-and trans-interactions would cause an increase in the mobility of PCP proteins remaining at the surface.

To determine how the stability of PCP proteins is modulated through the cell cycle, we performed fluorescence recovery after photobleaching (FRAP) analyses of Celsr1-3xGFP, Fz6-3xGFP and tdTomato-Vangl2 in basal epidermal cells during interphase and mitosis. Small regions with strong protein enrichment along vertically oriented (anterior-posterior) junctions of polarized cells, were selected for photobleaching and imaged continuously over a 3-minute recovery period (Fig. 2A-C). FRAP analysis of membrane-tdTomato expressed in the same cells as Celsr1-3xGFP was included as a control, as it is lipid anchored, uniformly localized and expected to freely diffuse within the plasma membrane (Fig. 2D)^30^. As expected, and consistent with FRAP experiments in *Drosophila*, the recovery curves for all three PCP components during interphase were characteristic of a relatively stable and immobile state, recovering more slowly and incompletely than membrane-tdTomato (Fig. 2’A-D’). By contrast, in mitosis, the fluorescence recovery curves for Celsr1-3xGFP, Fz6-3xGFP and tdTomato-Vangl2 plateaued at normalized intensities significantly above their respective plateaus in interphase (Fig. 2A’-C’), with significantly greater mobile fractions of all three proteins during mitosis than during interphase (Fig. 2E). Importantly, fluorescence recovery curves of membrane-tdTomato were similar across the two cell cycle phases (Fig. 2D’). Thus, an increase in lateral diffusivity is not a universal property of membrane associated proteins in mitosis. We conclude that upon entry to mitosis, Celsr1, Fz6 and Vangl2 transition from a stable state at polarized junctions during interphase to a more diffusive state within the membrane during mitosis.

**Figure 2.**
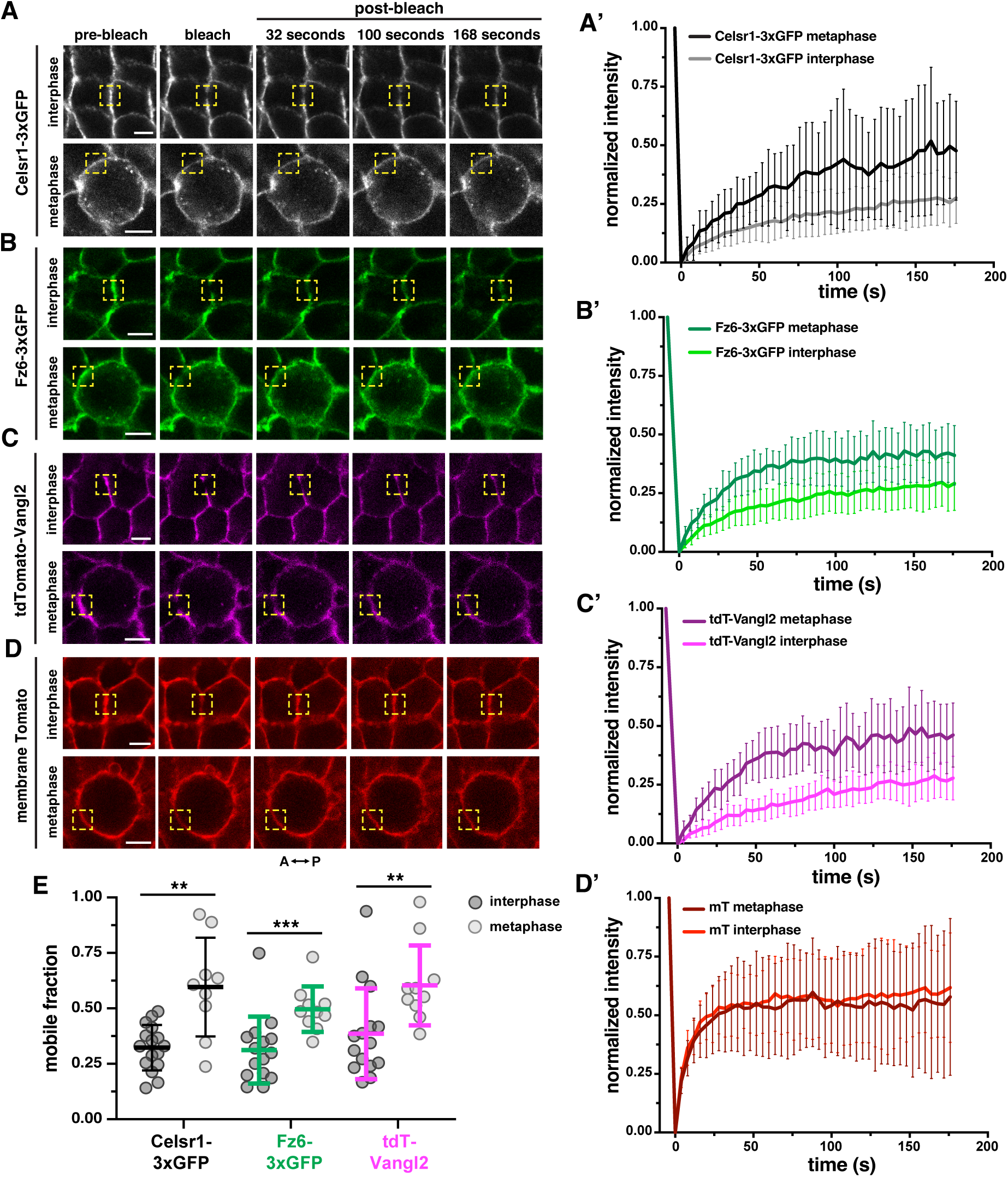
Membrane-localized PCP proteins are mobilized during mitosis. **(A-D)** Representative still frames of fluorescence recovery after photobleaching (FRAP) experiments at the membrane of *E*15.5 basal cells in interphase and metaphase. Bleached ROIs (1 μm diameter) are outlined by yellow dashed squares. Scale bar, 5 µm. (A) Celsr1-3xGFP, (B) Fz6-3xGFP, (C) tdT-Vangl2 and (D) membrane-tdTomato fluorescence are shown pre- and post-photobleaching. **(A’-D’)** Fluorescence recovery plots show the normalized mean intensity with standard deviations of bleach and recovery profiles plotted over time. The darker shade in each plot represents recovery during metaphase. The lighter shade represents recovery during interphase. (A’) Celsr1-3xGFP recovery curves, n = 21 ROIs for interphase and 15 ROIs for metaphase across 3 embryos. (B’) Fz6-3xGFP recovery curves, n = 15 ROIs for interphase and 15 ROIs for metaphase across 3 embryos. (C’) tdT-Vangl2 recovery curves, n = 15 ROIs for interphase and 15 ROIs for metaphase across 3 embryos. (D’) membrane Tomato recovery curves, n = 22 ROIs for interphase and 18 ROIs for metaphase across 3 embryos. **(E)** Mobile fractions of Celsr1-3xGFP (black), Fz6-3xGFP (green) and tdT-Vangl2 (magenta) during interphase (dark gray circles) and metaphase (light gray circles). Mann-Whitney test: **p = 0.0022, ***p = 0.0005, **p = 0.008. Bars indicate mean ± S.D. **(F)** Correlation of Fz6-3xGFP mobile fractions and tdT-Vangl2 mobile fractions during interphase (dark gray circles, solid black line) and metaphase (light gray circles, dashed red line). Correlations fit to simple linear regression y = mx + b, interphase: m = 1.191, b = 0.01387, metaphase: m = 1.293, b = −0.03788.

The increased mobility of PCP proteins observed in dividing cells suggests their localization is likely to become uniformly distributed in the membrane during mitosis. To test this directly, we visualized Fz6-3xGFP and tdT-Vangl2 localization in chimeric skins in which single, isolated Fz6-3xGFP; tdT-Vangl2 co-expressing cells were surrounded completely by unlabeled wild type cells. Strikingly, the mutually exclusive localization of Fz6-3xGFP and tdT-Vangl2 to opposite sides of the cell in interphase was lost as cells progressed through mitosis. As Fz6-3xGFP containing vesicles internalized from the posterior, tdT-Vangl2 spread evenly across the mitotic cell membrane losing its anterior enrichment (Fig. S2). Thus, mitotic internalization together with mobilization PCP proteins leads to a transient loss of planar polarity during mitosis.

### Increased Celsr1-3xGFP mobility during mitosis relies on Plk1

To investigate how PCP protein mobility is regulated during mitosis, we examined the role of the cell cycle kinase Polo-like kinase 1 (Plk1). Plk1 phosphorylates Celsr1 in mitosis, activating its endocytic motif and triggering its internalization^23^. Treatment of embryonic skin explants with Plk1 inhibitor (BI2536), arrests dividing cells in prometaphase and blocks Celsr1 mitotic internalization^23^. We therefore hypothesized that blocking Celsr1 endocytosis through Plk1 inhibition might immobilize Celsr1 at the plasma membrane. To test this, we performed FRAP analysis on *E*15.5 Celsr1-3xGFP skin explants treated with BI2536 or DMSO for a minimum of 3 hours (Fig. 3A-B). Live imaging confirmed that BI2536-treated cells remained arrested for at least 4 hours with visibly disorganized chromosomes compared to DMSO-treated controls (Movies S2 left and middle panels, Fig. S3) and, as expected, Celsr1-3xGFP was retained at the plasma membrane of prometaphase arrested cells (Fig. 3F). In FRAP experiments of junctional Celsr1-3xGFP, fluorescence recovered more slowly and less completely in dividing cells treated with BI2536 compared to DMSO controls. Fluorescence recovery curves of Plk1-inhibited cells arrested in mitosis were similar to cells in interphase (Fig. 3D). Moreover, the mobile fractions of Celsr1-3xGFP in interphase and BI2536-arrested cells were not significantly different (Fig. 3F). Only the mobile fraction of Celsr1-3xGFP in metaphase of control explants was significantly higher (Fig. 3F). We conclude that Plk1 kinase activity not only activates Celsr1 internalization but also promotes its mobilization within the plasma membrane.

**Figure 3.**
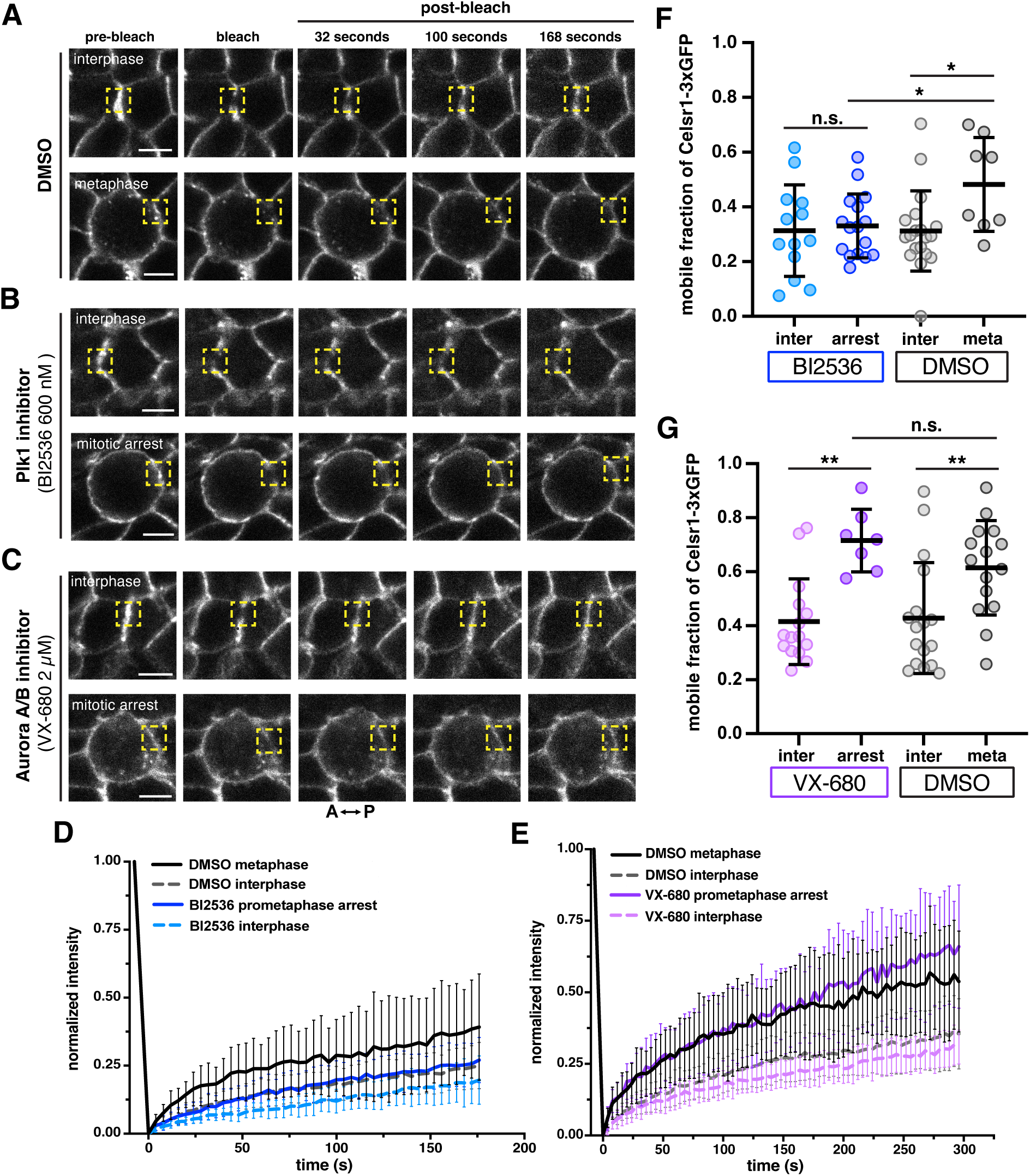
Increased Celsr1-3xGFP mobility during mitosis relies on Plk1. **(A-C)** Representative still frames from FRAP experiments at the membrane of *E*15.5 Celsr1-3xGFP expressing basal cells during interphase and either metaphase or arrest in prometaphase following exposure to **(A)** DMSO **(B)** Plk1 inhibitor (BI2536 600 nM, 3hr exposure) and **(C)** Aurora A/B inhibitor (VX-680 2 µM, 4.5 hr exposure). Bleached ROIs (1 μm diameter) are outlined by yellow dashed squares. Scale bars, 5 µm. **(D-E)** Fluorescence recovery plots show the normalized mean intensity with standard deviations of bleach and recovery profiles plotted over time. (D) Recovery curves of Celsr1-3xGFP during interphase (dashed light blue line) and prometaphase arrest (solid blue line) following BI2536 treatment, n = 19 ROIs for interphase and 17 ROIs for prometaphase arrest across 3 embryos. Recovery curves of Celsr1-3xGFP during interphase (dashed gray line) and metaphase (solid black line) following DMSO treatment, n = 19 ROIs for interphase and metaphase across 3 embryos. (E) Recovery curves of Celsr1-3xGFP during interphase (dashed light purple line) and prometaphase arrest (solid purple line) following VX-680 treatment, n = 17 ROIs for interphase and 19 ROIs for prometaphase arrest across 3 embryos. Recovery curves of Celsr1-3xGFP during interphase (dashed gray line) and metaphase (solid black line) following DMSO treatment, n = 18 ROIs for interphase and 16 ROIs for metaphase across 3 embryos. **(F)** Mobile fractions of Celsr1-3xGFP following BI2536 treatment during interphase (light blue circles) and prometaphase arrest (dark blue circles). Mobile fractions of Celsr1-3xGFP following DMSO treatment during interphase (light gray circles) and metaphase (dark gray circles). Mann-Whitney test: *p = 0.023, *p = 0.0112. **(G)** Mobile fractions of Celsr1-3xGFP following VX-680 treatment during interphase (light purple circles) and prometaphase arrest (dark purple circles). Mobile fractions of Celsr1-3xGFP following DMSO treatment during interphase (light gray circles) and metaphase (dark gray circles). Mann-Whitney test: **p = 0.0015, **p = 0.0061. Bars indicate mean ± S.D.

To distinguish whether mitotic arrest alone could account for the loss of Celsr1 mobility upon Plk1 inhibition, we treated Celsr1-3xGFP epidermal explants with an Aurora A/B inhibitor (VX-680), which arrests dividing cells in prometaphase but does not inhibit Celsr1 internalization (Movie S2 right panel, Fig. S3)^23^. Following 4.5 hours VX-680 treatment, we performed FRAP on Celsr1-3xGFP during interphase and mitosis and found their recovery curves to be similar to their control counterparts (Fig. 3C, E). The mobile fractions of Celsr1-3xGFP in control metaphase and VX-680 arrested cells were not significantly different, and both were higher than control and VX-680 treated cells in interphase (Fig. 3G). Thus, immobilization of Celsr1 upon Plk1 inhibition is not due to a change in overall membrane fluidity caused by mitotic arrest. Rather, we conclude that Celsr1’s mobilization within the membrane in mitosis is regulated through a Plk1-dependent mechanism. Whether Plk1 acts solely by promoting selective internalization of the stable Celsr1 pool or targets other factors that allow membrane-associated Celsr1 to diffuse along the membrane remains to be determined.

### Trafficking and inheritance of PCP-containing endosomes over the course of cell division

Thus far we have shown that membrane-associated PCP proteins switch from an immobile state in interphase to a diffusive state in mitosis. Together with the internalization of PCP proteins from the cell surface, these changes in PCP protein dynamics effectively erase planar polarization during mitosis. To investigate the mechanisms that restore polarity following mitosis, we hypothesized that internalized PCP proteins are transported equally to both daughters and delivered to the membrane in a polarized manner. To test this, we followed PCP-containing vesicles over the course of cell division and quantified vesicle numbers at anterior and posterior positions during mitosis and cytokinesis. Consistent with Celsr1’s localization to both sides of the junction in interphase, roughly equal numbers of Celsr1-3xGFP-containing endosomes formed on the anterior and posterior halves of cells in mitosis (Fig. 4A). Endosomes containing Fz6-3xGFP, by contrast, were positioned with a bias to the anterior half, while vesicles with tdT-Vangl2, though far fewer in number, were mainly present on the posterior half (Fig. 4A). Thus, Fz6 and Vangl2 containing endosomes enter the cytoplasm unequally in mitosis.

**Figure 4.**
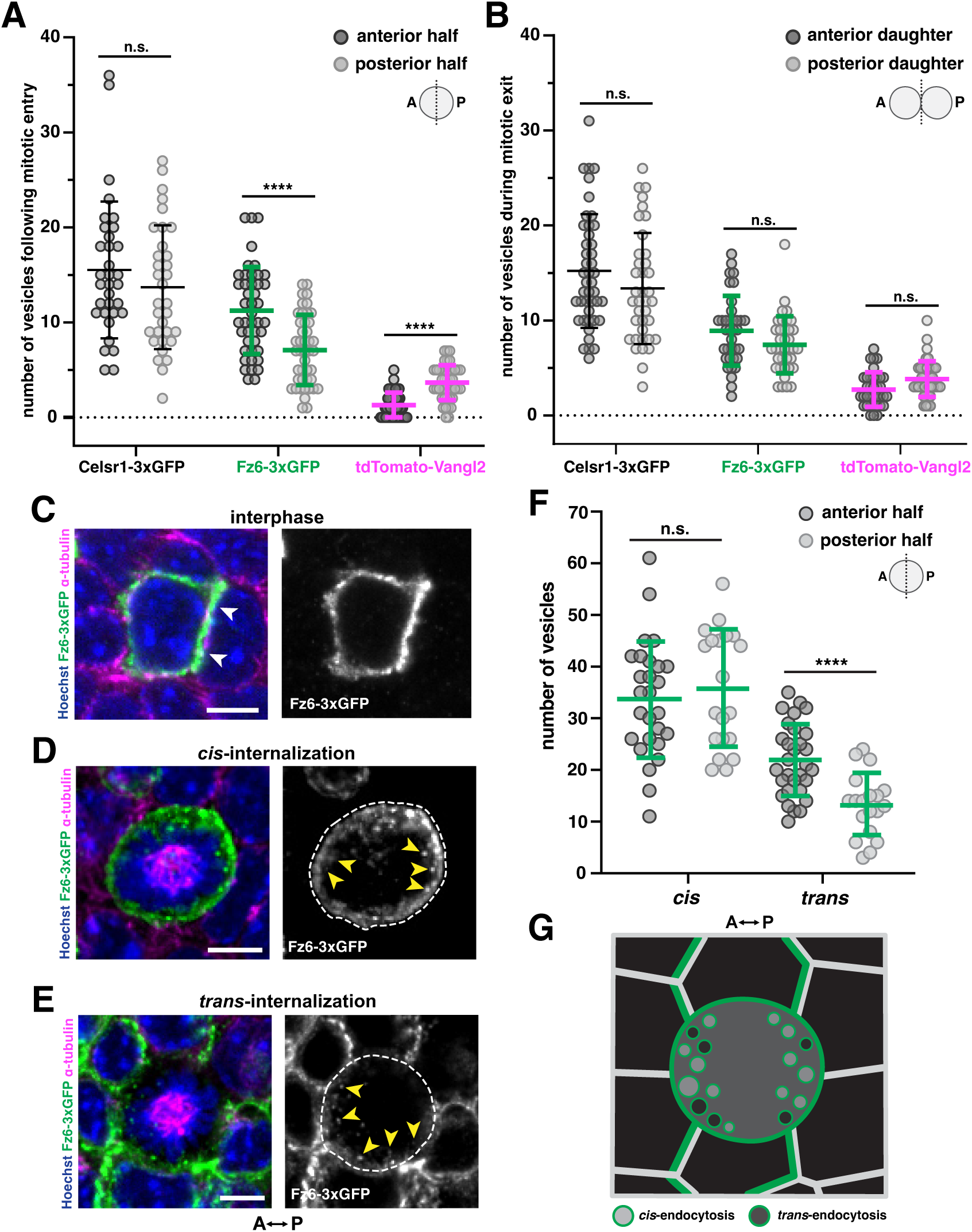
Trafficking and inheritance of PCP-containing endosomes over the course of cell division. **(A)** Quantification of Celsr1-3xGFP, Fz6-3xGFP and tdT-Vangl2-containing endosomes in the anterior and posterior halves of dividing basal cells in prophase to metaphase, n = 30 mitotic cells expressing Celsr1-3xGFP across 6 embryos; 42 mitotic cells expressing Fz6-3xGFP across 4 embryos; 44 mitotic cells expressing tdT-Vangl2 across 4 embryos. Unpaired two-tailed t-test: ****p < 0.0001. Bars indicate mean ± S.D. **(B)** Quantification of Celsr1-3xGFP, Fz6-3xGFP and tdT-Vangl2-containing endosomes in anterior and posterior daughter cells, n = 39 daughter pairs expressing Celsr1-3xGFP across 6 embryos; 36 daughter pairs expressing Fz6-3xGFP across 4 embryos; 36 daughter pairs expressing tdT-Vangl2 across 4 embryos. **(C-E)** Representative images of basal cells from *E*15.5 *WT : Fz6-3xGFP* chimeric skin with nuclei labeled by Hoechst (blue) and the microtubule cytoskeleton labeled by α-tubulin (magenta). (C) Example of a Fz6-3xGFP cell in interphase surrounded by unlabeled *WT* cells. Arrowheads (white) point to the asymmetric localization of Fz6-3xGFP (green) enriched at the posterior cell border. (D) Example of a mitotic Fz6-3xGFP cell (outlined by white dashed line) surrounded by *WT* cells. *Cis*-internalization of Fz6-3xGFP at the anterior and posterior halves are marked by yellow arrowheads. (E) Example of a *WT* cell in mitosis (outlined by white dashed line) surrounded by Fz6-3xGFP cells. *Trans*-internalization of Fz6-3xGFP at the anterior and posterior halves marked by yellow arrowheads. **(F)** Distribution of Fz6-3XGFP *cis-* and *trans-*endosomes in anterior (darker circles) and posterior (lighter circles) halves of mitotic cells; n = 27 Fz6-3xGFP mitotic cells with anterior *WT* neighbors (*cis-*anterior); 20 Fz6-3xGFP mitotic cells with posterior *WT* neighbors across 3 embryos (*cis-*posterior); 30 *WT* mitotic cells with anterior Fz6-3xGFP (*trans-*anterior); 19 *WT* mitotic cells with posterior Fz6-3xGFP neighbors (*trans-*posterior); all across 3 embryos, Mann-Whitney test: ****p < 0.0001. Scale bars, 5 µm. **(G)** Diagram illustrating the anterior bias in accumulation of Fz6-3xGFP vesicles derived from *trans-*endocytosis, whereas *cis-*endocytosed vesicles accumulate fairly evenly on both sides of the cell.

Given their localization patterns in interphase (Fz6 on the posterior, Vangl2 on the anterior), the asymmetry of Fz6 and Vangl2 endosomes in dividing cells appears opposite the expected distribution (compare Fig. 1A to Fig. 4A). However, in a previous study, we found that the majority of Vangl2-containing vesicles in dividing cells accumulate posteriorly and arise from trans-endocytosis of Celsr1-Vangl2 complexes from neighbors ^27^. Therefore, the large number of anterior Fz6-3xGFP vesicles in dividing cells might also arise via transendocytosis of Fz6 protein from the posterior side of their immediate neighbor. To test this, we generated chimeric mouse embryos consisting of a mix of Fz6-3xGFP and WT cells and examined the distribution of Fz6-3xGFP endosomes in dividing cells at the interfaces of unlabeled and GFP-positive cells (Fig. 4C). This allowed us to distinguish vesicles that internalized from the membrane of the dividing cell itself (cis-internalization) from those that entered the dividing cell from its neighbors (trans-internalization) (Fig. 4D, E). Endosomes arising from cis-internalization accumulated on both anterior and posterior sides of dividing cells in roughly equal numbers (Fig. 4D, F), whereas Fz6-3xGFP vesicles deriving from trans-endocytosis were greater on the anterior (Fig. 4E, F). Together, the sum of *cis-* and *trans-*internalized vesicles can account for the overall greater number of Fz6-3xGFP vesicles that accumulate on the dividing cell’s anterior side, despite Fz6’s posterior enrichment in interphase (Fig. 4G).

We next compared the number of Celsr1-3xGFP, Fz6-3xGFP and tdTomato-Vangl2-containing endosomes between anterior and posterior daughter cells in cytokinesis and found, consistent with previous findings in fixed skin samples, all three PCP proteins were equally inherited in both daughter cells (Fig. 4B)^22^. Therefore, any biases in internalization at mitotic entry are corrected during mitotic exit through the equal partitioning of endosomes in daughter cells.

### PCP-containing endosomes are not delivered in a polarized manner to restore asymmetry following mitosis

Endosomes are known to be delivered to specific cellular locations by polarized delivery along microtubules. In the *Drosophila* wing epithelium, directed transport of Flamingo (Fmi), Frizzled (Fz) and Disheveled (Dsh) along aligned noncentrosomal microtubules is thought to bias their asymmetry during polarity establishment^31–34^. Therefore, we assessed whether PCP protein-containing endosomes are trafficked via microtubule-based transport in mitosis. Using stimulated emission depleted (STED) super-resolution microscopy, we visualized the localization of Celsr1-3xGFP-containing endosomes with respect to the microtubule cytoskeleton at various stages of the cell cycle (Fig. 5A, Fig. S2A). In interphase, Celsr1-3xGFP was localized along anterior-posterior cell junctions and was only minimally associated with the microtubule cytoskeleton (Fig. 5A, Fig. S4A, far left panels). In metaphase, Celsr1-3xGFP-containing endosomes were abundant in the cytoplasm, but these were not associated with spindle or astral microtubules (Fig. 5A, Fig. S4A, middle left panels). In telophase and cytokinesis, some Celsr1-3xGFP-containing endosomes were associated with midzone microtubules (Fig. 5A, Fig. S4A middle right and right panels), suggesting a possible mechanism for transport between daughter cells. However, at all stages of cell division, the vast majority of Celsr1-3xGFP-containing endosomes were not associated with microtubules but were dispersed throughout the cytoplasm. Similarly, we observed that only a small fraction of Fz6-3xGFP-containing endosomes associated with spindle or midzone microtubules (Fig. S3B). Together these data suggest that although some Celsr1-3xGFP and Fz6-3xGFP may be transported along microtubules in mitosis, their partitioning and redelivery to the plasma membrane cannot rely solely on this method of transport.

**Figure 5.**
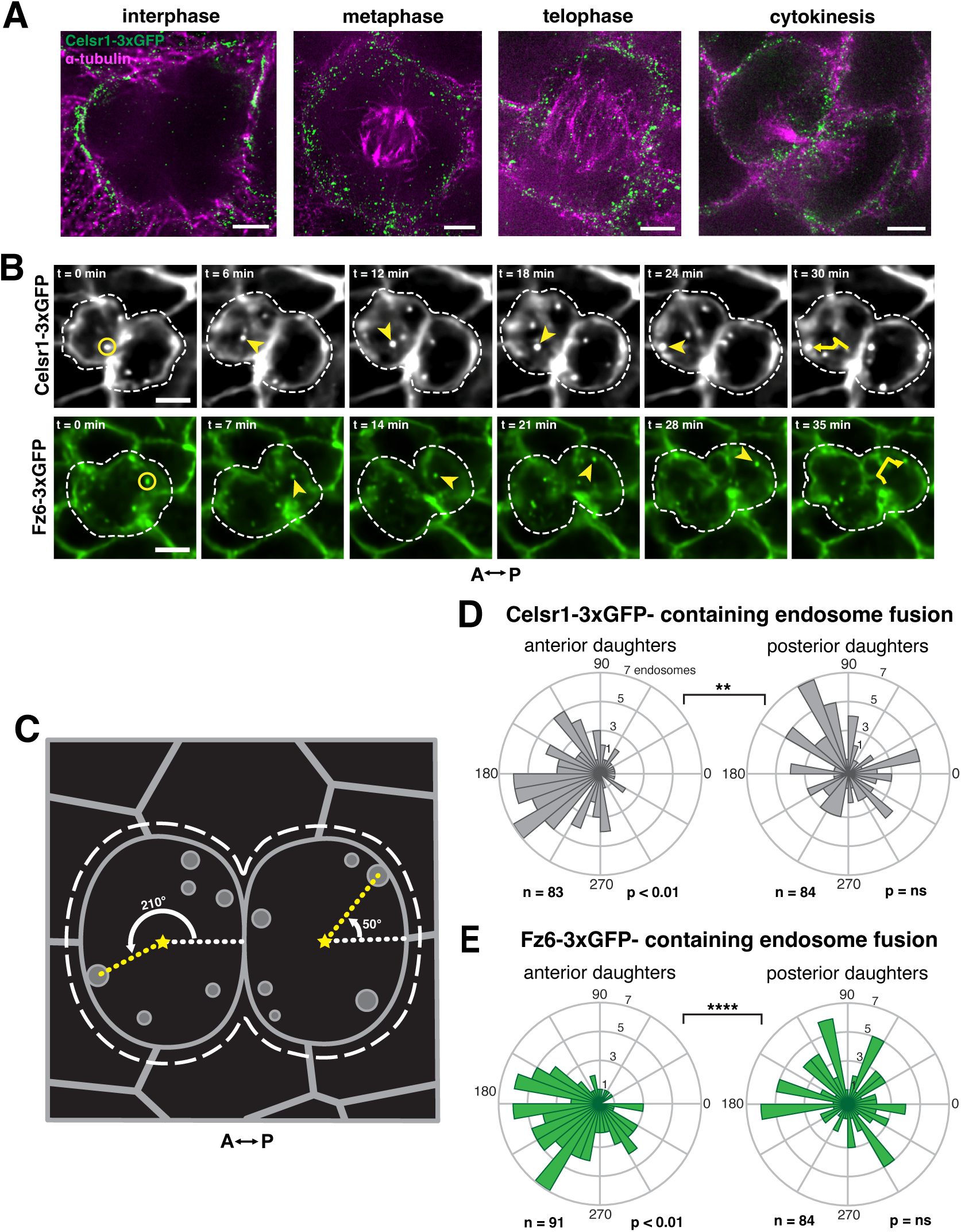
PCP-containing endosomes are not delivered in a polarized manner to restore asymmetry following mitosis. **(A)** Stimulated emission depleted microscopy (STED) super resolution imaging of Celsr1-3xGFP containing endosomes and mitotic spindle microtubules. Whole mount of *E*15.5 *Celsr1-3xGFP* skins were labeled for GFP (green) and α-tubulin (magenta). Representative images of basal cells in interphase, metaphase, telophase and cytokinesis showing minimal colocalization of Celsr1-3xGFP-containing endosomes with the microtubule cytoskeleton. Scale bar, 2.5 µm. **(B)** Still frames from representative time-lapse of Celsr1-3xGFP- and Fz6-3xGFP-containing endosomes (shown in grayscale and green, respectively) fusing to the plasma membrane of daughter cells during cytokinesis. White dashed lines demarcate daughter cells. At t=0 min, yellow circles mark the Celsr1-3xGFP- and Fz6-3xGFP -containing endosomes that eventually fuse to the plasma membrane in the last timepoint. Yellow arrowheads point to the endosome in the direction from the previous timepoint. The trajectories are shown with a yellow arrow in the last timepoint. Scale bar, 5 µm. **(C)** Diagram illustrating how angles of endosome fusion were calculated. Yellow stars represent the centroids of each daughter cell. **(D-E)** Rose plots showing the angle distribution of Celsr1-3xGFP (gray) and Fz6-3xGFP-containing (green) endosomes fusing to the plasma membrane in anterior and posterior daughters. (D) Celsr1-3xGFP-containing endosome fusion angle distribution, n = 83 Celsr1-3xGFP-containing endosomes in anterior daughters across 7 embryos; 84 Celsr1-3xGFP-containing endosomes in posterior daughters across 8 embryos. One-sample Kuiper’s Test: *p < 0.01 (E) Fz6-3xGFP-containing endosome fusion angle distribution, n = 91 Fz6-3xGFP-containing endosomes in anterior daughters across 7 embryos; 84 Fz6-3xGFP-containing endosomes in posterior daughters across 6 embryos. One-sample Kuiper’s Test: *p < 0.01; Fz6-3xGFP expressing posterior daughter, p>0.05. Two-sample Kuiper’s Test: **p = 0.0036, ****p < 0.0001.

Next, we asked whether vesicles carrying PCP proteins fuse asymmetrically to the plasma membrane to restore their asymmetric localization. Given their respective distributions during interphase, we expected that fusion of Celsr1-3xGFP-containing endosomes would be biased toward anterior and posterior cell borders of daughter cells, whereas Fz6-3xGFP-containing endosomes would fuse preferentially to posterior cell borders. Using live imaging, we tracked Celsr1-3xGFP- and Fz6-3xGFP-containing endosomes during cytokinesis and observed where they fuse to the plasma membrane (Fig. 5B). Given their sparse numbers, we chose to forgo analysis of tdT-Vangl2-containing endosomes. To quantify the positions of endosome fusion to the plasma membrane, we measured the angle between each daughter cell’s centroid and the point of endosome fusion (Fig. 5C). Regardless of the orientation in which cells divided along the A-P axis, both Celsr1-3xGFP- and Fz6-3xGFP-containing endosomes fused with a bias to the anterior cell border of anterior daughter cells (Fig. 5D, E). In posterior daughters, fusion positions were random and unbiased (Fig. 5D, E). Thus, PCP-containing endosomes are not delivered to ‘correct’ positions in cytokinesis and, at least in anterior daughters, Fz6 is delivered to the opposite side of where it accumulates in interphase. Thus, redelivery of Celsr1-3xGFP and Fz6-3xGFP to the plasma membrane does not restore their asymmetric localization. Further refinement of their distributions is required following membrane localization.

### Membrane-associated PCP proteins remain highly mobile through cytokinesis

Asymmetric PCP protein localization is eventually restored in daughter cells, but this does not occur through polarized fusion of PCP protein-containing endosomes in cytokinesis. How then are PCP proteins able to assume their asymmetric localizations following division? If PCP proteins are able to laterally diffuse within the plasma membrane following endosome fusion, cis- and trans-interactions with other PCP proteins could immobilize and stabilize PCP complexes through time, ultimately restoring their polarized distributions. To quantify the lateral mobility of PCP proteins during cell cycle exit, we performed FRAP analysis of Celsr1-3xGFP and Fz6-3xGFP;tdT-Vangl2 expressing cells undergoing cytokinesis (Fig. 6A). We found that junctional Celsr1-3xGFP, Fz6-3xGFP and tdTomato-Vangl2 fluorescence recovered faster and more completely during cytokinesis than during interphase (Fig. 6B-D, Fig. S5). The mobile fractions of all three PCP components were significantly higher during cytokinesis than during interphase (Fig. 6F). By contrast, fluorescence recovery of membrane-tdTomato (mT) was unchanged between interphase and cytokinesis, indicating the increased mobility of PCP proteins is not due to a broad increase in membrane fluidity (Fig. 6E). Together our data show that following endosome fusion, transmembrane PCP components remain diffusive within the membrane during cytokinesis to later become stabilized in interphase.

**Figure 6.**
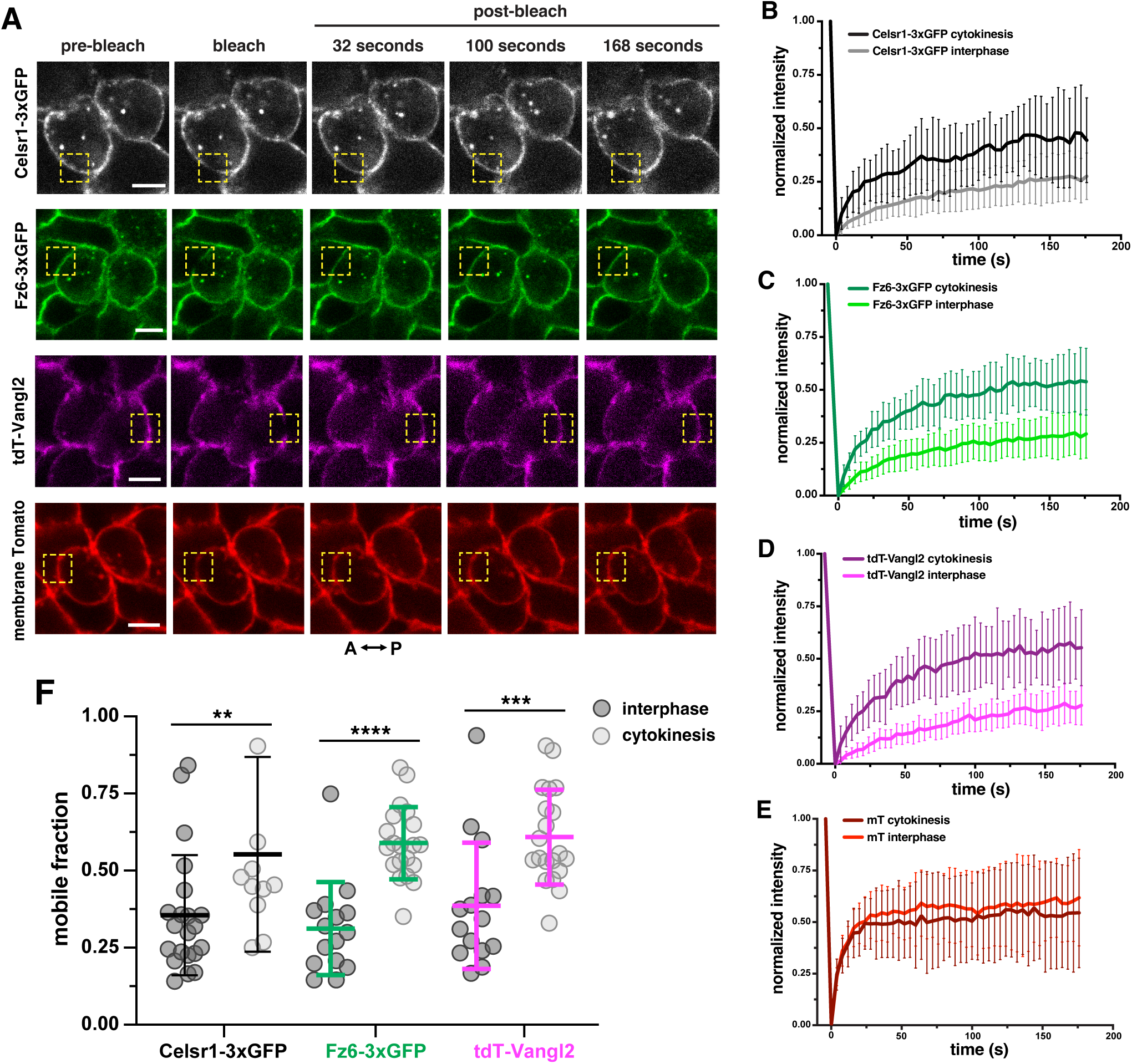
Membrane-associated PCP proteins remain highly mobile through cytokinesis. **(A)** Representative still frames of FRAP analysis of Celsr1-3xGFP (grayscale), Fz6-3xGFP (green), tdT-Vangl2 (magenta) or membrane Tomato (red) at the plasma membrane of basal cells in cytokinesis. Bleach ROIs (1 μm diameter) are outlined by yellow dashed squares. Scale bars, 5 µm. **(B-E)** Fluorescence recovery plots show the normalized mean intensity with standard deviations of bleach and recovery profiles plotted over time. The darker line in each plot represents recovery during cytokinesis and the lighter line represents recovery during interphase. (B) Celsr1-3xGFP recovery curves, n = 21 ROIs for interphase and 34 ROIs for cytokinesis across 3 embryos. (C) Fz6-3xGFP recovery curves, n = 15 ROIs for interphase and 28 ROIs for cytokinesis across 3 embryos. (D) tdT-Vangl2 recovery curves, n = 15 ROIs for interphase and 28 ROIs for cytokinesis across 3 embryos. (E) membrane Tomato recovery curves, n = 22 ROIs for interphase and 34 ROIs for cytokinesis across 3 embryos. **(F)** Mobile fractions of Celsr1-3xGFP (black), Fz6-3xGFP (green) and tdT-Vangl2 (magenta) during interphase (dark gray circles) and cytokinesis (light gray circles). Mann-Whitney test: **p = 0.0066, ****p < 0.0001, ***p = 0.0004. Bars indicate mean ± S.D. **(G)** Correlation of Fz6-3xGFP and tdT-Vangl2 mobile fractions during interphase (dark gray circles, solid black line) and cytokinesis (light gray circles, dashed black line). Correlations fit to simple linear regression (y = mx + b), interphase: m = 1.191, b = 0.01387, cytokinesis: m = 0.6936, b = 0.1996.

Given that PCP proteins remain highly mobile within the membrane during cytokinesis, their asymmetric localization is unlikely to be restored until the following interphase. To test this, we returned to *Fz6-3XGFP; tdT-Vangl2 : WT* chimeric skins and identified dual-reporter expressing cells in telophase and cytokinesis that were surrounded by unlabeled neighbors. Consistent with their diffusivity at these stages, Fz6 and Vangl2 were uniformly distributed around the surface of cells in telophase and early stages of cytokinesis. Only during late stages of cytokinesis did their enrichment along anterior and posterior cell edges begin to emerge (Fig. S6).

### PCP is restored non-cell autonomously

Thus far we have shown that PCP proteins undergo a shift toward higher mobility upon mitotic entry that persists through cytokinesis. To transition back to a stable state at anterior and posterior junctions in interphase, PCP proteins might be captured and stabilized through trans-interactions with their neighbors. We therefore hypothesized that Celsr1 in neighboring cells could stabilize post-mitotic PCP components non-cell autonomously. To test this, we used embryo aggregation to generate chimeric mouse embryos consisting of a mix of Fz6-3xGFP;tdT-Vangl2 expressing cells and either unlabeled wild type or *Celsr1-/-* mutant cells (Fig. 7A)^25^. Due to the difficulties in performing live imaging and FRAP analysis of such rare chimeric embryos, we used clustering as a proxy for stabilization. PCP proteins organize and cluster into discrete membrane subdomains, referred to as puncta^24, 26, 28, 29^, whose assembly is promoted by Fmi/Celsr1, which participates in both homotypic trans-adhesive and cis-clustering interactions^17, 26, 28^. To test the role of Celsr1 in Fz6 and Vangl2 clustering, we identified dual PCP reporter-expressing cells that directly neighbored unlabeled WT or *Celsr1-/-* cells and characterized Fz6-3xGFP and tdT-Vangl2 clusters along shared edges by measuring: 1) the size of clusters and 2) the fractional area of the cell edge occupied by clusters. Per each cell, puncta were identified by first performing thresholding in the GFP and RFP channels. The median threshold value was then calculated and subsequently applied to the anterior and posterior edges of cells.

**Figure 7.**
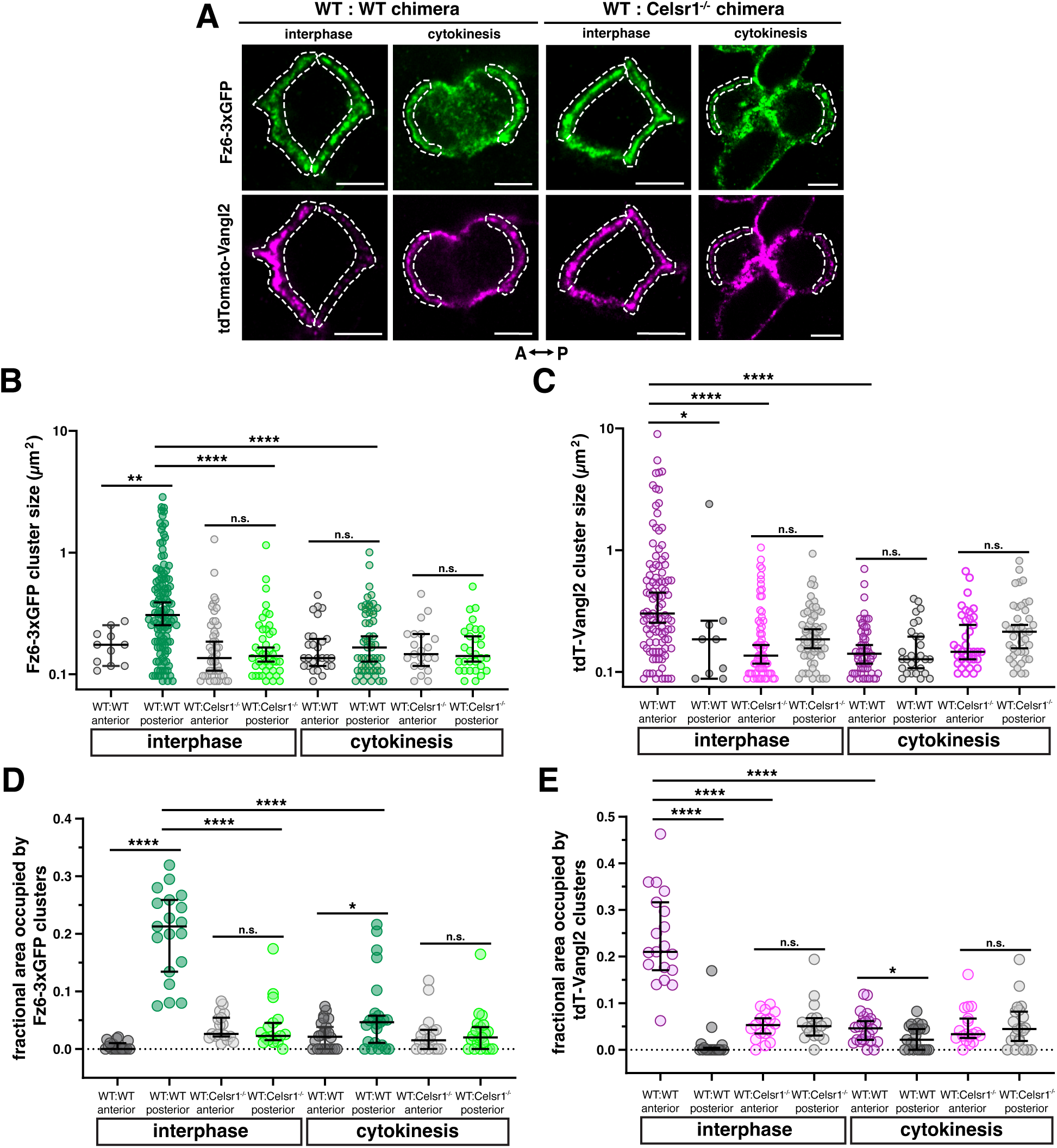
PCP is restored non-cell autonomously. **(A)** Representative images of Fz6-3xGFP (green) and tdT-Vangl2 (magenta) in interphase and cytokinesis taken from *E*15.5 *Fz6-3xGFP;tdT-Vangl2 : WT (WT* : *WT)* and *Fz6-3xGFP;tdT-Vangl2 : Celsr1-/-* (*WT* : *Celsr1-/-*) chimeric skins. Fz6-3xGFP- and tdT-Vangl2 expressing cells are surrounded either entirely or partially by unlabeled WT or Celsr1-/- cells. Only anterior and posterior edges shared between fluorescent and non-fluorescent cells (white dashed lines) were analyzed. Scale bars, 5 µm. **(B)** Quantification of Fz6-3xGFP cluster size, n = 7 anterior *WT* : *WT* interphase junctions; 19 posterior *WT* : *WT* interphase junctions; 17 anterior *WT* : *Celsr1-/-* interphase junctions; 16 posterior *WT* : *Celsr1-/-* interphase junctions; 14 anterior *WT* : *WT* cytokinesis interfaces; 18 posterior *WT* : *WT* cytokinesis interfaces; 12 anterior *WT* : *Celsr1-/-* cytokinesis interfaces; 15 posterior *WT* : *Celsr1-/-* cytokinesis interfaces; all across 3 embryos, Mann-Whitney test: **p = 0.0037, ****p < 0.0001. **(C)** Quantification of tdT-Vangl2 cluster size, n = 19 anterior WT : WT interphase junctions; 5posterior *WT* : *WT* interphase junctions; 16 anterior *WT* : *Celsr1-/-* interphase junctions; 16 posterior *WT* : *Celsr1-/-* interphase junctions; 22 anterior *WT* : *WT* cytokinesis interfaces; 14 posterior *WT* : *WT* cytokinesis interfaces; 17 anterior *WT* : *Celsr1-/-* cytokinesis interfaces; 16 posterior *WT* : *Celsr1-/-* cytokinesis interfaces; all across 3 embryos, Mann-Whitney test: *p = 0.0269, ****p < 0.0001. Bars indicate mean ± S.D. Note: only one data point met cluster criteria for posterior WT : WT interphase junction. **(D)** Quantification of the fractional area occupied by Fz6-3xGFP clusters, n = 19 anterior and 19 posterior WT : WT interphase junctions; 17 anterior and 17 posterior *WT* : *Celsr1-/-* interphase junctions; 24 anterior and 23 posterior *WT* : *WT* cytokinesis interfaces; 18 anterior and 21 posterior *WT* : *Celsr1-/-*cytokinesis interfaces; all across 3 embryos, Mann-Whitney test: ****p < 0.0001, *p = 0.0497. Bars indicate mean ± S.D. **(E)** Quantification of the fractional area occupied by tdT-Vangl2 clusters, n = 19 anterior and 19 posterior *WT* : *WT* interphase junctions; 17 anterior and 17 posterior *WT* : *Celsr1-/-* interphase junctions; 24 anterior and 23 posterior *WT* : *WT* cytokinesis interfaces; 18 anterior and 21 posterior *WT* : *Celsr1-/-* cytokinesis interfaces; all across 3 embryos, Mann-Whitney test: ****p < 0.0001, *p = 0.0216. Bars indicate median with 95% Confidence Interval.

To first determine the cluster characteristics of Fz6-3xGFP and tdT-Vangl2 in polarized cells in interphase, we compared puncta distributions along anterior and posterior junctions of *Fz6-3xGFP;tdT-Vangl2 : WT* (WT : WT) chimeric interfaces. Fz6-3xGFP clusters at the posterior were larger and occupied more area compared to anterior junctions (Fig. 7B, D). Conversely, tdT-Vangl2 clusters were found almost exclusively on anterior junctions, and compared to posterior tdT-Vangl2 clusters, clusters at the anterior were on average larger in size and occupied more area along the edge than posterior clusters (Fig. 7C, E). Thus, consistent with their low mobility at posterior and anterior junctions, Fz6-3xGFP and tdT-Vangl2 are arranged into clustered assemblies during interphTo test whether Celsr1 is required non-autonomously for Fz6-3xGFP and tdT-Vangl2 stabilization during interphase, we measured clustering at *Fz6-3xGFP;tdT-Vangl2 : Celsr1-/-* interfaces (WT : Celsr1-/-). When Celsr1 is absent in neighbors, Fz6-3xGFP clusters along posterior junctions were significantly smaller and occupied less area along the junction compared to when Celsr1 is present (Fig. 7B, D). Similarly, tdT-Vangl2 clusters along WT : Celsr1-/- anterior junctions were significantly smaller and occupied less area along the junction compared to their WT : WT anterior counterparts (Fig. 7C, E). Celsr1 is therefore required to non-autonomously cluster and stabilize Fz6-3xGFP and tdT-Vangl2 during interphase.

Next we measured cluster characteristics in cells undergoing cytokinesis and found at WT : WT edges that Fz6-3XGFP and tdT-Vangl2 puncta were smaller and occupied less area along their respective posterior and anterior edges compared to their counterparts in interphase (Fig. 7B-E). These data correlate well with the higher mobility of Fz6 and Vangl2 observed during cytokinesis and suggest that only when daughter cells have entered interphase do Fz6-3xGFP and tdT-Vangl2 assemble into larger, stabilized complexes. Nevertheless, when comparing anterior and posterior edges of daughter cells in cytokinesis, small, but significant differences in Fz6-3XGFP and tdT-Vangl2 clustering were observed. Fz6-3xGFP clusters occupied a larger fractional area along posterior edges, whereas tdT-Vangl2 puncta were more prominent along the anterior sides (Fig. 7D, E). These anterior-posterior differences observed in cytokinesis were lost when Celsr1 was lacking in neighbors (Fig. 7D, E). Altogether, these data underscore the requirement of Celsr1 to non-cell autonomously stabilize Fz6-3xGFP and tdT-Vangl2 post-mitosis. However, rather than Fz6 and Vangl2 immediately immobilizing at the end of mitosis, stabilization likely occurs through a gradual process involving Celsr1 trans-interactions and cis-clustering as daughter cells are reincorporated into the epithelium.

## DISCUSSION

The asymmetric localization of polarity proteins is established through dynamic regulation of polarity protein ‘states’, for example, immobile versus mobile, bound versus unbound, and clustered versus diffuse^9^. A mobile state allows polarity proteins to move to new locations in the cell whereas immobility retains asymmetric localizations. Components of the planar polarity complex are known to exist in different states of mobility and clustering over the course of PCP establishment^26, 28, 35–40^, but given the long time scales over which PCP asymmetry emerges, for example, many hours in *Drosophila* and days in mouse epidermis, deciphering how PCP protein dynamics are regulated over the course of polarization is a challenge. By investigating how PCP proteins are dynamically remodeled during mitosis, we have monitored their state transitions over minutes rather than days. This revealed that the mobility of PCP components greatly increases in mitosis, both through internalization into cytoplasmic vesicles, as we have previously described^22, 23, 27^, and through increasing lateral diffusion in the membrane. Endocytosis of junctional PCP proteins likely contributes to the mobility of PCP proteins that remain at the membrane by reducing the net stabilizing *cis-* and *trans-*interactions between the remaining intercellular PCP complexes. This increased mobility persists during cytokinesis, even after PCP-containing vesicles are recycled to the surface, allowing PCP proteins to laterally diffuse and eventually be captured and stabilized at AP interfaces by Celsr1-containing complexes within their neighbors. Importantly, oscillations in PCP protein mobility are controlled by the mitotic kinase Plk1, thereby coupling PCP dynamics with cell cycle progression.

We considered the possibility that directed transport of PCP-containing endosomes might restore asymmetry after division. Exocytic vesicles containing Fmi, Fz and Dvl transit along aligned microtubules in the *Drosophila* wing, which is thought to bias global PCP orientation^31–34^. Furthermore, asymmetric endosome transport along central spindle microtubules determines asymmetric cell fate in *Drosophila* sensory organ precursors^41^. However, we do not observe directed transport of PCP-containing endosomes along spindle microtubules in dividing epidermal cells. Given that the division planes of basal cells do not consistently align with the PCP axis^22, 42^, a mechanism whereby endosomes travel along spindle microtubules, would not necessarily restore asymmetry in the correct orientation. Instead, we find that Fz6-containing endosomes fuse with a slight bias toward the anterior side of the cell, opposite its eventual accumulation in interphase. One possible explanation for this anterior bias is the retention of anteriorly localized Vangl2^27^, which may target Celsr1- and Fz6-containing endosomes to the anterior cell border. Nevertheless, vesicular transport and fusion cannot explain how PCP asymmetry is restored following division. Rather, an increase in lateral diffusion within the membrane appears to be the main mechanism for moving PCP proteins to their eventual destination in cytokinesis.

In multiple polarity systems, establishment and/or maintenance of asymmetry is associated with the assembly of polarity complexes into micron-scale aggregates that display different dynamics compared to non-clustered pools^43–49^. In the *Drosophila* wing, PCP complexes organize into immobile puncta along cell junctions, and during polarity establishment, mobility inversely correlates with the degree of asymmetry^28, 29, 37, 40, 48^. In mouse epidermal cells, PCP complexes are also organized into puncta, in part through cis-adhesive interactions mediated by the atypical cadherin Celsr1^24, 26^. Extracellular tethering though Celsr1 trans-binding also restricts PCP protein mobility^26^, but surprisingly, Celsr1 adhesive interactions contribute only weakly to PCP complex clustering and stabilization during cytokinesis. Only later, during interphase, do PCP proteins assemble into clustered and immobile junctional complexes. This suggests that Celsr1 adhesive interactions might be inhibited when cells divide, allowing for polarity to be restored gradually as daughters are reintegrated into the epithelium. Interestingly, *Drosophila* Fmi is proposed to exist in open and closed forms^21^, and perhaps Celsr1 is maintained in a closed confirmation until mitosis is complete.

The switching of polarity protein states in coordination with the cell cycle appears to be a general feature of polarized cells, allowing for cells to adjust their polarity for stage-specific functions. For example, apicobasal polarity proteins in some epithelial tissues are transiently released from the epithelial cell cortex during mitosis, regaining their cortical localization following division^10, 50^. These oscillations in cell polarity are thought to allow repositioning of the cell adhesion machinery between daughter cells as they are reincorporated into the epithelium, or to allow basolateral polarity proteins to associate with the spindle orientation machinery. Mechanistically, oscillations in apicobasal polarity and PAR complex behavior have been linked to Cdk1, AurK or Plk1 activity, thereby closely linking polarity dynamics and state transitions with mitotic progression^50–56^. For the planar polarity complex, Plk1 phosphorylates S/T resides surrounding Celsr1’s dileucine endocytic motif, activating its removal from the membrane via clathrin-mediated endocytosis^23^. Here we find that Plk1 activity also promotes the mobility of PCP proteins within the membrane, although we cannot distinguish if increased mobility is solely a secondary consequence of internalization. However, the persistence Fz6 and Vangl2 mobility in cytokinesis, even after PCP components have returned to the surface, suggests that Plk1 activity could contribute directly to a mobile state.

We propose that the changes to planar polarity organization and mobility in mitosis allow the orientation of polarity to be adjusted and realigned in coordination with epithelial growth. Epidermal progenitors proliferative rapidly during embryonic development and yet, despite the extensive reorganization of cellular architecture that accompanies division, the global orientation of planar polarity remains aligned with the body axes^22, 24^. Intuitively, mitosis should be disruptive to polarity alignment, but in a recent computational model of planar polarity establishment, the introduction of divisions in which polarity is erased and restored enhanced long-range alignment of polarity across the epithelium (Thayambath and Belmonte, 2025). Simulations of planar polarity establishment in which a small tissue proliferates displays better polarity alignment than a tissue that starts large (Thayambath and Belmonte, 2025). Our data demonstrate mechanistically how this can be achieved – through increased lateral mobility of PCP complexes within the membrane and eventual capture and stabilization through trans-interactions with neighbors. Thus, when mitosis is coupled to an increase in PCP complex mobility, it may not hinder planar polarity but rather facilitate its global order.

## ACKNOWLEDGEMENTS

We thank Gary Laevsky and Sha Wang of the Confocal Imaging Center at Princeton, a Nikon Center of Excellence for assistance and advice with all imaging experiments, Katherine Little for assistance with mouse husbandry and genotyping, members of the Devenport lab past and present, Jean Schwarzbauer and Ricardo Mallarino for helpful discussions, and Rishabh Sharan for assistance with data quantitation. This work was supported by grants from NIH NIAMS R01AR066070 and R01AR068320 (to D.D), NSF grant IOS2052511 (to D.D), NIH NIAMS F31AR077407 (to L.B), NJCCR Postdoctoral Award COCR24PDF018 (to P.S.), NIH NIGMS T32GM007388 (to D.O. and L.B.)

## MATERIALS AND METHODS

**Table 1.**
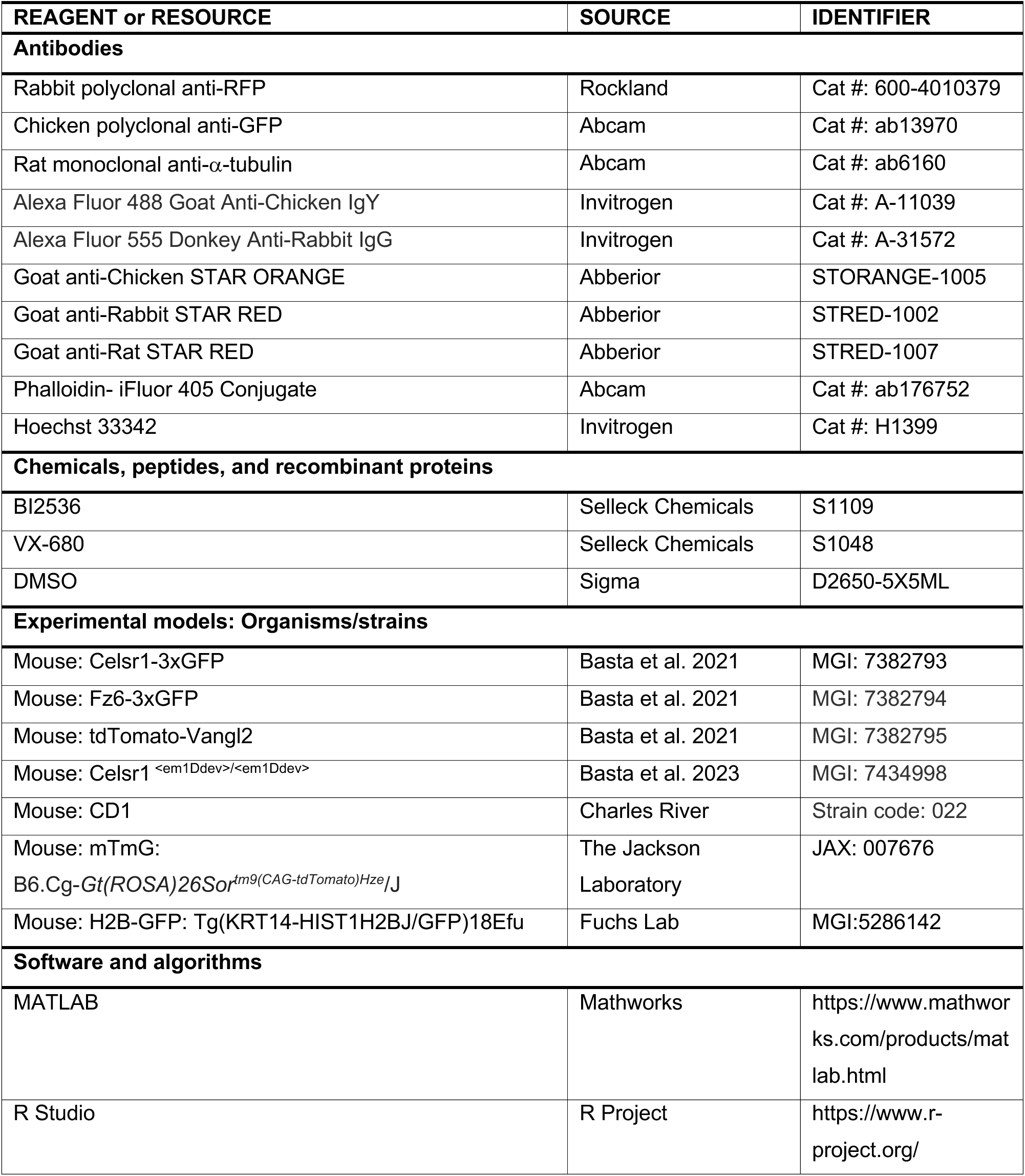
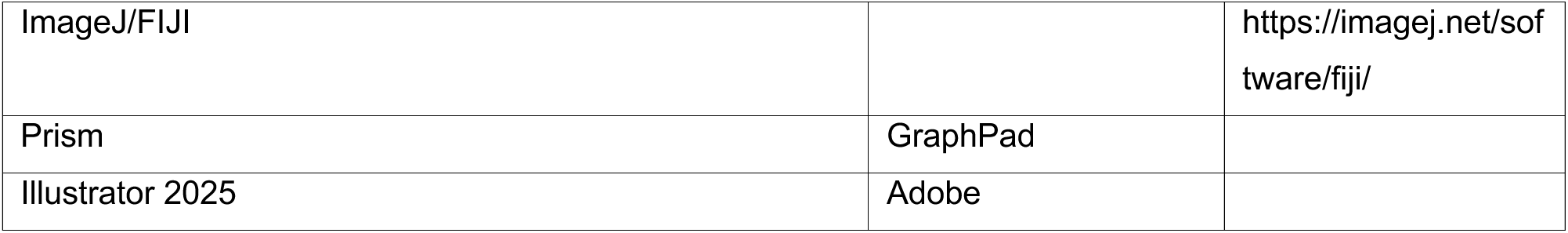
Key resources.

### Mouse lines and breeding

Mice were housed following the Guide for the Care and Use of Laboratory Animals in an AAALAC-accredited facility. Animal maintenance and husbandry were done in accordance with the Animal Welfare Act. All animal procedures were approved by Princeton University’s Institutional Animal Care and Use Committee (IACUC). Experiments were performed on *E*15.5 embryos obtained by mating *Celsr1-3xGFP; mTmG* homozygous mice, *Fz6-3xGFP* homozygous mice, *Fz6-3xGFP; tdTomato-Vangl2* mice maintained as heterozygous for the *tdTomato-Vangl2* allele^24^ and H2B-GFP; mTmG homozygous mice^30, 57^.

### Production of Chimeras

Chimeric *WT : Fz6-3xGFP;tdTomato-Vangl2*, *Fz6-3xGFP;tdTomato-Vangl2 : Celsr1^<em1Ddev>/<em1Ddev>^* and *WT : Fz6-3xGFP E*15.5 embryos were produced through aggregation of previously cryopreserved embryos. Embryos were cryopreserved by rapid cooling as described in Kuleshova and Shaw, 2000^58^. Briefly 2-cell embryos were collected at *E*1.5 by flushing the oviduct of plugged females. Embryos were collected and cryopreserved in glass capillary straws filled with cryopreservation solution and plunged in liquid nitrogen. To thaw, embryo straws were removed from liquid nitrogen using cooled forceps and warmed in order to clear the cryopreservation solution. Embryos were then washed and cultured overnight at 37°C, 6% CO_2_ and 5% O_2_ until they reached the *E*2.5 morula stage the next day.

Morula stage embryos were used to create aggregation chimeras using protocols described in Manipulating the mouse embryo, 2014 (Specifically chapter 12 – Production of Chimeras Protocols, 8 and 10). Briefly the zona pellucida was removed from morula stage embryos and washed before being moved to a previously prepared aggregation plate. One morula from each genotype was carefully placed into each aggregation well so that the two morulas can aggregate into a chimera. Embryos were cultured overnight and assessed for aggregation and development into a blastocyst. Successfully aggregated chimeras that had developed into blastocysts (now stage E3.5) were transferred into the uterus of 2.5 days post coitum pseudo-pregnant CD1 females and dissected 12 days later at E15.5. Embryos were genotyped following dissection.

### Immunofluorescence and fixed sample imaging

For fixing and staining of embryonic epidermis, *E*15.5 embryos were dissected in PBS and fixed in 4% paraformaldehyde for 1 hour at room temperature. Backskins were blocked overnight at 4°C in PBT2 (PBS with 0.2% Triton X-100) with 2% normal goat serum, 2% normal donkey serum, 1% bovine serum albumin (BSA) and 1% fish gelatin. Samples were then incubated in primary antibodies in PBT2 block at 4°C overnight. Following primary incubation, samples were washed thoroughly in PBT2 at room temperature and incubated in secondary antibodies in PBT2 overnight at 4°C or at room temperature for two hours. Following another round of washes in PBT2 and a final wash in PBS, skins were mounted in Prolong Gold (Molecular Probes) and stored at −20°C.

The primary antibodies used were: rabbit anti-RFP (1:200, Rockland, 600-4010379); chicken anti-GFP (1:2000, Abcam, ab6556); and rat anti-alpha-tubulin (1:500 Abcam, ab6160). The secondary antibodies used were: Alexa Fluor-488 and −555 (1:2000; Invitrogen, A-11039 and A-31572) plus Hoechst (1:2000, Thermo Fisher, H1399) or Phalloidin-iFluor 405 (1:2000, Abcam, ab176752). The secondary antibodies used for STED imaging were: goat anti-chicken STAR ORANGE (1:100, Abberior, STORANGE-1005), goat anti-Rabbit STAR RED (1:100, Abberior, STRED-1002) and goat anti-rat STAR RED (1:100, Abberior, STRED-1007).

Images were acquired on a Nikon A1R-HD25 confocal microscope coupled to an inverted Ti-2 stand using a Plan Apo 100/1.49 NA oil-immersion objective (Nikon). STED images were also acquired using the Plan Apo 100/1.49 NA oil-immersion objective on the same confocal microscope with a STEDYCON stimulated emission depletion (STED) module. Nikon NIS-Elements and ImageJ were used for image processing. NIS-Elements AR analysis was used to deconvolve STED images before additional processing with ImageJ.

### Live imaging and movie processing

Live imaging of *E*15.5 epidermal explants was performed as previously described^59^. *E*15.5 dorsal flank skin explants were dissected in PBS and mounted dermal side down onto an agar pad (1% agarose gel with F-media containing 10% fetal bovine serum). The agar pads were mounted on a 35-mm air permeable Lummox membrane dish (Sarstedt) with the epidermal side of the skin in contact with the membrane. Images were acquired using a Plan Apo 60/1.4 NA oil immersion objective on a Nikon Ti-2 inverted CSU-W1 SoRa Spinning Disk microscope equipped with a Tokai-Hit stage top incubator and an environmental incubation chamber, both maintained at 37° and 5% CO_2_. Z stacks with 0.5 mm z steps were acquired every 2-3 minutes for 2-3 hours. Cells within hair follicle placodes and at the edge of the explant were not imaged to avoid differences in planar cell polarity that could occur as a result of morphogenesis or wound healing response, respectively. NIS-Elements AR analysis and ImageJ were used for denoising and processing images.

### Quantification of vesicle distribution, fusion analysis, and clustering analysis

#### Quantification of PCP-protein containing vesicles during division

To analyze the number of PCP-protein containing vesicles following mitotic entry, prometaphase/metaphase cells from live imaging were bisected in half using ImageJ. Anterior and posterior halves were assigned relative to the AP axis. Vesicles in both halves of the cells were counted manually from the highest apical plane of the cell where vesicles were visible, to the lowest basal plane. Similarly, following cytokinesis, daughter cells were assigned as either anterior or posterior relative to the AP axis and PCP-containing vesicles were manually counted in both daughter cells.

#### Analysis of Fz6-3xGFP cis- and trans-endocytosis. E15.5 Fz6-3xGFP

*WT* chimeric backskins were first dissected, fixed and stained as described above. To quantify the distribution of endosome resulting from Fz6-3xGFP *cis-* or *trans-*endocytosis, cells in prometaphase/metaphase either expressing Fz6-3xGFP and surrounded/flanked by unlabeled WT cells (*cis*-endocytosis) or not expressing Fz6-3xGFP (unlabeled WT) and surrounded/flanked by Fz6-3xGFP expressing cells (*trans*-endocytosis) were bisected in half using ImageJ. Vesicles were counted manually at the anterior and posterior sides of the mitotic cell as described above.

#### Fusion analysis

Vesicles that fused to the plasma membrane during cytokinesis were first identified from live imaging movies. To quantify the position of endosome fusion to the plasma membrane, we used ImageJ to first calculate the centroids of daughter cells. The angle between each of the daughter cell’s centroid and the point of endosome fusion was then measured. As cells were oriented relative to the AP axis, 0° represented the posterior side of a cell and 180° represented the anterior. The distribution of vesicle fusion was plotted in MATLAB. Significance was performed in R Studio and calculated using the kuiper.test from the circular package and the kuiper.2samp function from the kuiper.2samp package for one- and two-sample Kuiper’s tests, respectively.

#### Fz6-3xGFP and tdTomato-Vangl2 cluster analysis in chimeras

*E*15.5 *WT : Fz6-3xGFP; tdTomato-Vangl2 (WT : WT)*, and *Fz6-3xGFP; tdTomato-Vangl2 : Celsr1^<em1Ddev>/<em1Ddev>^*(WT : Celsr1^-/-^) chimeric backskins were fixed, stained, and imaged using confocal as described above. As a note, tdTomato-Vangl2 was either heterozygous or homozygous, however we were unable to confirm due to limitations from genotyping in chimeras. Isolated double positive cells (GFP- and tdTomato-positive, termed double positive) in interphase completely surrounded by non-expressing *WT* or *Celsr1^-/-^*neighbors were identified. Given the sparsity of double-positive cells in cytokinesis surrounded entirely by non-expressing *WT or Celsr1^-/-^*cells, we also imaged those that were partially flanked by cells expressing GFP and tdTomato. In these cases, to ensure that we were only quantifying clusters contributed by one cell, we only quantified the double-positive edges shared with non-expressing WT or Celsr1^-/-^ edges.

Using ImageJ, background pixel values for WT : WT and WT : Celsr1^-/-^ samples were calculated from 20 ROIs along junctions non-expressing WT and 20 ROIs along non-expressing Celsr1^-/-^ junctions. The average background pixel value for non-expressing WT and Celsr1^-/-^ cells was then calculated and subtracted from corresponding WT : WT or WT : Celsr1^-/-^ images. The entire cell membrane of cells in interphase and cytokinesis (excluding cleavage furrow) were isolated, followed by thresholding using the Otsu pre-set (with further adjustments to the min and max if needed for reliable recapitulation of the junction) on both the Fz6-3xGFP and tdTomato-Vangl2 channels to create a mask of protein expression along the membrane. A median + 3.5 median absolute deviation threshold value was then calculated for each channel using MATLAB to produce a final threshold value. For interphase cells, we isolated the anterior and posterior edges of double-positive cells that neighbored non-expressing cells. Likewise, for cells in cytokinesis, we isolated the anterior edges of anterior daughters and the posterior edges of posterior daughters that bordered non-expressing cells. Using the final threshold value, we isolated clusters that were larger than 9 pixel^2^ (to eliminate clusters that were not Nyquist sampled by at least three pixels in any dimension, assuming a square shape) and calculated the size and area coverage of clusters as well as the area of cell edge. Cluster size, cluster density (# of clusters/total area of edge) and fractional area occupied by clusters (total area occupied by clusters/total area of edge) were calculated in Microsoft Excel. Data and significance were plotted using Graphpad Prism.

### FRAP and inhibitor experiments

#### FRAP of transmembrane PCP proteins across cell cycle stages

*E*15.5 dorsal flank skin explants were dissected and mounted for live imaging as described above. Explants were imaged using a Plan Apo 100/1.49 NA oil-immersion objective (with additional zoom that rendered the pixel size 110 nm) on a Nikon A1R-HD25 confocal microscope equipped with a Tokai-Hit stage top incubation chamber maintained at 37° and 5% CO_2_. Magnification, laser power (for bleach and acquisition), pixel dwell time, and acquisition rate were all kept constant across all measurements. The FRAP acquisition sequence consisted of three reference pre-bleach images followed by bleach to a 1-μm diameter circular ROI and subsequent acquisition at 4 sec intervals across 45 frames to capture fluorescence recovery. For interphase recovery measurements, bleach ROIs were placed at anterior-posterior polarized cell junctions. For metaphase, bleach ROIs were placed at the plasma membrane on either the anterior or posterior sides of the mitotic cell (since no significant difference in mobility was observed at this stage). For cytokinesis, bleach ROIs were placed at the plasma membrane on either the anterior side of anterior daughters or the posterior side of posterior daughters (to avoid the newly forming shared junction between daughter cells). Background pixel values were calculated from three ROIs outside each explant and subtracted from each time series. Images in each timeseries were checked for any Z-drift and corrected for XY-drift in NIS-Elements. Three reference ROIs were made along non-bleached junctions of each movie to correct for overall bleaching during image acquisition. Each series was background and bleach corrected. The ROI values were extracted from NIS-Elements and analyzed in Microsoft Excel and GraphPad Prism. The corrected intensity profiles were then normalized as *(F_t_–F_bleach_)/(F_ini_–F_bleach_)*, where, F_t_ is intensity of the ROI at any time point, F_bleach_ is intensity at time point immediately after bleaching, and F_ini_ is mean ROI intensity of pre-bleach frames. To quantify the mobile fractions, each recovery curve was fitted to the exponential one phase association equation in Graphpad Prism. The fitted Plateau and Y_0_ values from the mean trace were used to determine the averaged *mobile fraction* = *(Plateau-Y_0_)/(1- Y_0_)*. Each recovery curve was fitted to the model with an r-squared value >0.70 to determine the individual mobile fractions using the above equation. All plots were created in Graphpad Prism.

#### FRAP experiments in the presence of cell cycle inhibitors

For Plk1 inhibition, agar pads for explant growth were supplemented with either 600 nM BI2536 (Selleck Chemicals S1109) dissolved in DMSO or the equivalent volume of DMSO (Sigma D2650-5X5ML). Explants were incubated at 37° and 5% CO_2_ in a cell culture incubator for a minimum of 3 hours before FRAP was performed. For Aurora A/B inhibition, agar pads were supplemented with either 2 μM VX-680 (Selleck Chemicals S1048) dissolved in DMSO or the equivalent volume of DMSO. Explants were incubated at 37° and 5% CO_2_ for a minimum of 4.5 hours before FRAP was performed. Images were acquired using the same microscope set up as described above. Magnification, laser power, pixel dwell time and acquisition rate were kept the same as the above FRAP experiments. Following bleach, recovery images were acquired at 4 second intervals over 75 frames. Three background images were taken outside each DMSO-control and inhibitor-treated explant with background values calculated using ImageJ. Subsequent analysis was performed on each drift corrected movie using NIS-Elements and ImageJ as described above. Recovery curves and mobile fractions were calculated and plotted using GraphPad Prism as described above.

### Statistical Analyses

Statistical analyses were performed as indicated in each Fig. legend. Unpaired t-test was calculated using Excel to determine the statistical significance in vesicle distribution following mitotic entry and exit. Mann-Whitney test and one-phase association was calculated using Graphpad Prism. One-sample and two-sample Kuiper’s tests were used for circular statistics and calculated using R Studio.

**Table 2.** Mouse genotypes.

| **Fig.** | **Genotypes** |
| --- | --- |
| **1B** | *+/+ : Fz6-3xGFP/Fz6-3xGFP; tdTomato-Vangl2/tdTomato-Vangl2*<br>*+/+ : Fz6-3xGFP/Fz6-3xGFP; tdTomato-Vangl2/+* |
| **1D** | *Celsr1-3xGFP/Celsr1-3xGFP*<br>*Fz6-3xGFP/Fz6-3xGFP; tdTomato-Vangl2/ tdTomato-Vangl2*<br>*Fz6-3xGFP/Fz6-3xGFP; tdT-Vangl2/+* |
| **2A, D** | *Celsr1-3xGFP/Celsr1-3xGFP; mTmG/mTmG* |
| **2B, C** | *Fz6-3xGFP/ Fz6-3xGFP; tdTomato-Vangl2/ tdTomato-Vangl2*<br>*Fz6-3xGFP/Fz6-3xGFP; tdT-Vangl2/+* |
| **3A, D** | *Celsr1-3xGFP/Celsr1-3xGFP* |
| **4C-E** | *+/+ : Fz6-3xGFP/Fz6-3xGFP* |
| **5A** | *Celsr1-3xGFP/Celsr1-3xGFP* |
| **5B** | *Celsr1-3xGFP/Celsr1-3xGFP*<br>*Fz6-3xGFP/ Fz6-3xGFP; tdTomato-Vangl2/ tdTomato-Vangl2*<br>*Fz6-3xGFP/Fz6-3xGFP; tdT-Vangl2/+* |
| **6A** | *Celsr1-3xGFP/Celsr1-3xGFP; mTmG/mTmG*<br>*Fz6-3xGFP/ Fz6-3xGFP; tdTomato-Vangl2/ tdTomato-Vangl2*<br>*Fz6-3xGFP/Fz6-3xGFP; tdT-Vangl2/+* |
| **7A** | *+/+; Fz6-3xGFP/Fz6-3xGFP; tdTomato-Vangl2/tdTomato-Vangl2*<br>*+/+; Fz6-3xGFP/Fz6-3xGFP; tdT-Vangl2/+* |
| **S1** | *Celsr1-3xGFP/Celsr1-3xGFP*<br>*Fz6-3xGFP/Fz6-3xGFP; tdTomato-Vangl2/tdTomato-Vangl2*<br>*Fz6-3xGFP/Fz6-3xGFP; tdT-Vangl2/+* |
| **S3A** | *Celsr1-3xGFP/Celsr1-3xGFP* |
| **S3B** | *Fz6-3xGFP/Fz6-3xGFP* |
| **Movie S1** | *K14-H2B-GFP/K14-H2B-GFP; mTmG/mTmG* |
| **Movie S2** | *Celsr1-3xGFP/Celsr1-3xGFP*<br>*Fz6-3xGFP/Fz6-3xGFP; tdTomato-Vangl2/tdTomato-Vangl2*<br>*Fz6-3xGFP/Fz6-3xGFP; tdT-Vangl2/+* |

\* Genotypes with tdTomato-Vangl2 were either heterozygous or homozygous, given that there were limitations from genotyping.

## Notes

### Competing Interest Statement

The authors have declared no competing interest.

### Summary of Updates

Author list has been updated such they are now listed in the correct order.

