## Supplemental Figures for "Cell cycle-dependent transitions in PCP protein mobility erase and restore planar cell polarity in mitosis"

### Supplementary Figure S1

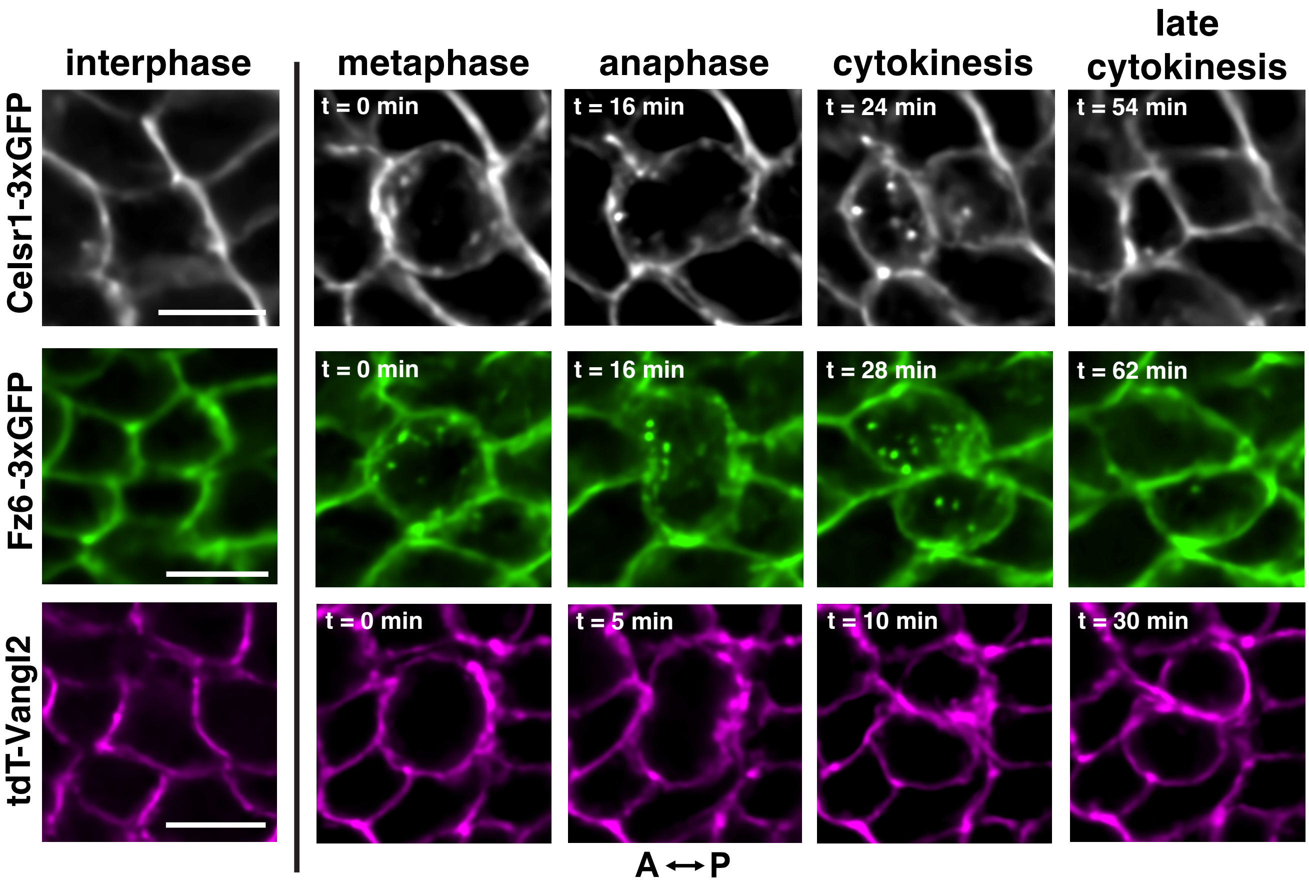


**Supplementary Figure S1. Still frames from live-imaging of mitotic basal cells in E15.5 PCP reporter epidermis.**
Additional examples of still frames from live imaging of *E*15.5 basal cells in *Celsr1-3xGFP* (top panels) and *Fz6-3xGFP;tdT-Vangl2* (middle and bottom panels) skin explants. Distribution of Celsr1-3xGFP (grayscale), Fz6-3xGFP (green) and tdT-Vangl2 (magenta) at interphase and from metaphase through late cytokinesis. Scale bars, 10 $\mu$m.

### Supplementary Figure S2

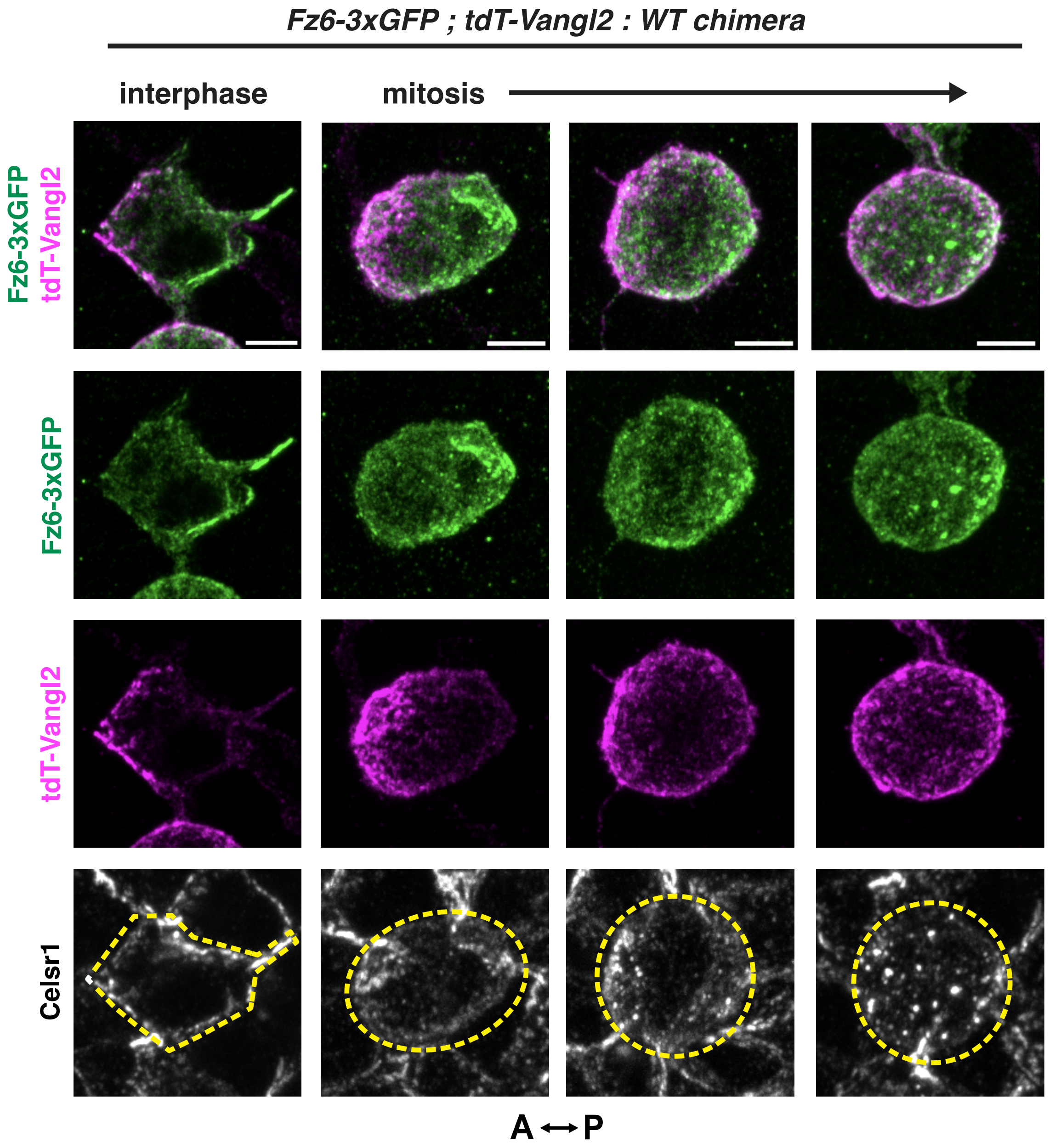


**Supplementary Figure S2. Asymmetric localization of PCP proteins is lost during mitosis.** Representative examples of isolated basal epidermal cells co-expressing Fz6-3xGFP (green) and tdT-Vangl2 (magenta) that are completely surrounded by unlabeled wild-type cells in *E*15.5 chimeric mouse embryos (*Fz6-3xGFP;tdT-Vangl2* : *WT*). Chimeric skins are also labeled with Celsr1 antibodies (grayscale). Shown are maximum intensity projections of cells in interphase and progressively later stages of mitosis (indicated by the extent of Celsr1 internalization). Note the progressive loss of anterior-localized Vangl2 and posterior-localized Fz6 at the cell surface as cells progress through mitosis. Scale bars, 5 $\mu$m.

### Supplementary Figure S3

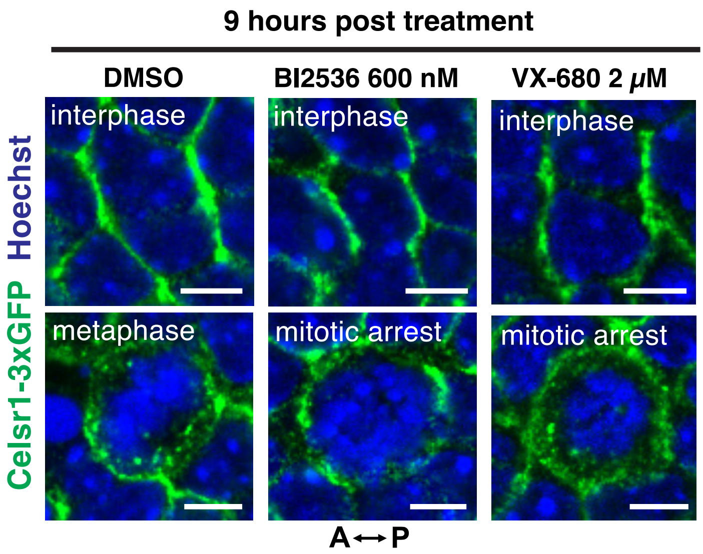


**Supplementary Figure S3. Basal cells arrest in prometaphase following 9 hours of Plk1 and Aurora A/B inhibitor treatment.** Representative examples of basal cells from *E*15.5 *Celsr1-3xGFP* skin explants (green) 9 hours post treatment with DMSO, BI2536 and VX-680. Basal cells in interphase (top panels) and metaphase or prometaphase arrest (bottom panels) are labeled with Hoechst (blue). Celsr1-3xGFP is internalized during mitosis and following prometaphase arrest with Aurora A/B inhibition, however it remains retained at the plasma membrane following Plk1 inhibition. Scale bars, 5 $\mu$m.

### Supplementary Figure S4

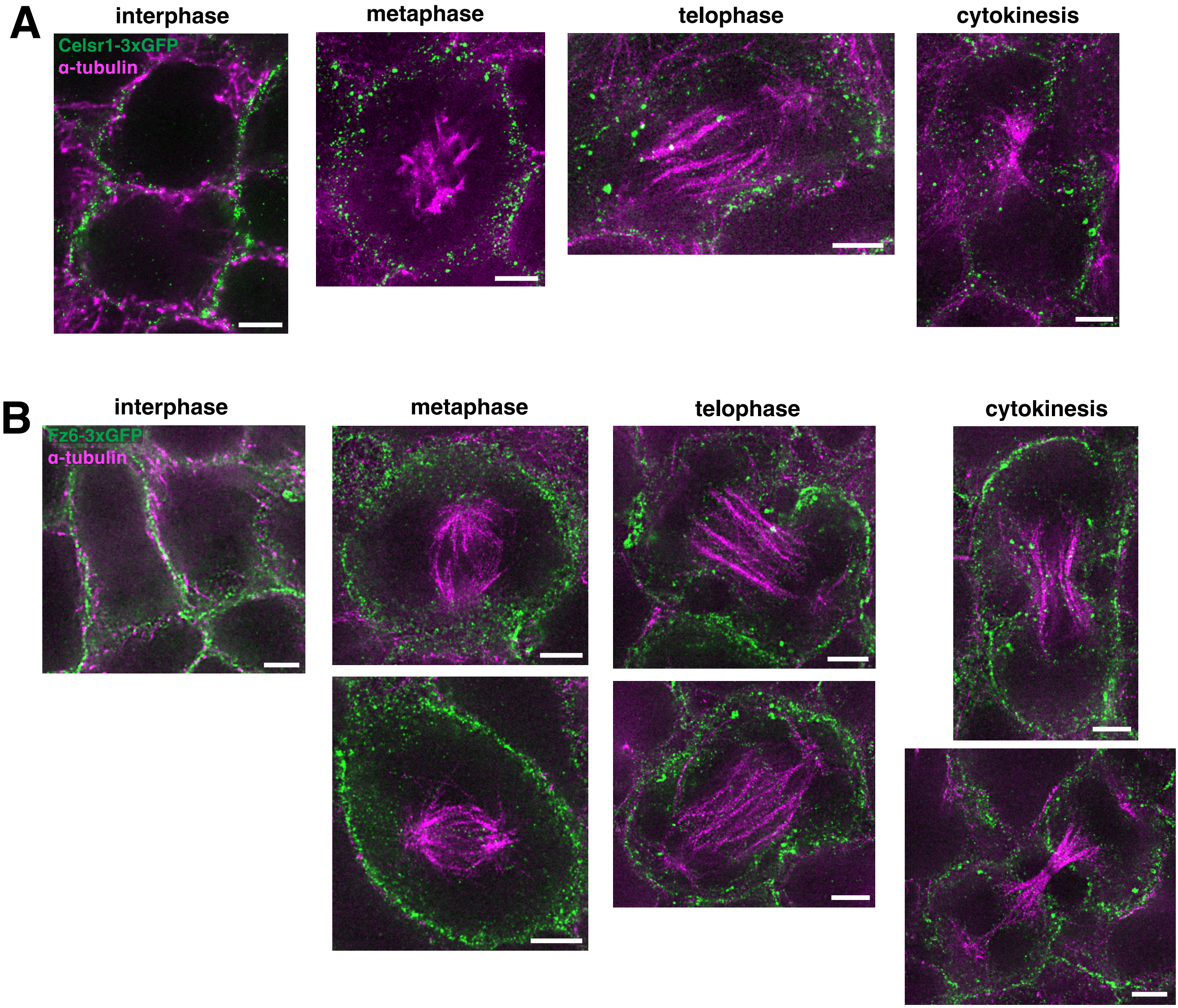


**Supplementary Figure S4. Celsr1-3xGFP and Fz6-3xGFP minimally colocalize with the microtubule cytoskeleton during division.** (A-B) STED super resolution imaging of Celsr1-3xGFP and Fz6-3xGFP containing endosomes and mitotic spindle microtubules. Whole mount *E*15.5 (A) *Celsr1-3xGFP* and (B) *Fz6-3xGFP* skins were labeled for GFP (green) and -tubulin (magenta) and imaged using STED microscopy. Representative images of basal cells in interphase, metaphase, telophase and cytokinesis showing minimal colocalization of Celsr1-3xGFP- and Fz6-3XGFP-containing endosomes with the microtubule cytoskeleton. Scale bars, 2.5 $\mu$m.

### Supplementary Figure S5

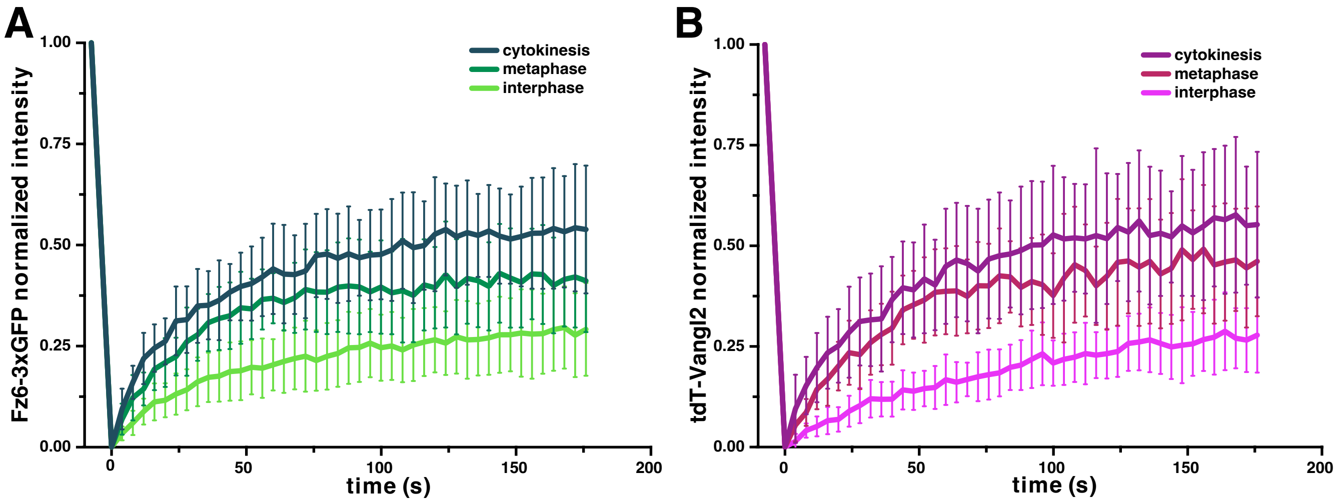


**Supplementary Figure S5. FRAP fluorescence recovery curves of Fz6-3xGFP and tdT-Vangl2 during interphase, metaphase and cytokinesis.**

(A) Fz6-3xGFP normalized mean intensities and standard deviations for bleach and fluorescence recoveries during interphase (light green), metaphase (dark green) and cytokinesis (dark teal) plotted over time. Fz6-3xGFP recovery curves, n = 15 ROIs for interphase, 15 ROIs for metaphase and 28 ROIs for cytokinesis across 3 embryos. (B) tdT-Vangl2 normalized mean intensities and standard deviations for bleach and fluorescence recoveries during interphase (magenta), metaphase (maroon) and cytokinesis (dark purple) plotted over time. tdT-Vangl2 recovery curves, n = 15 ROIs for interphase and 15 ROIs for metaphase and 28 ROIs for cytokinesis across 3 embryos.

### Supplementary Figure S6

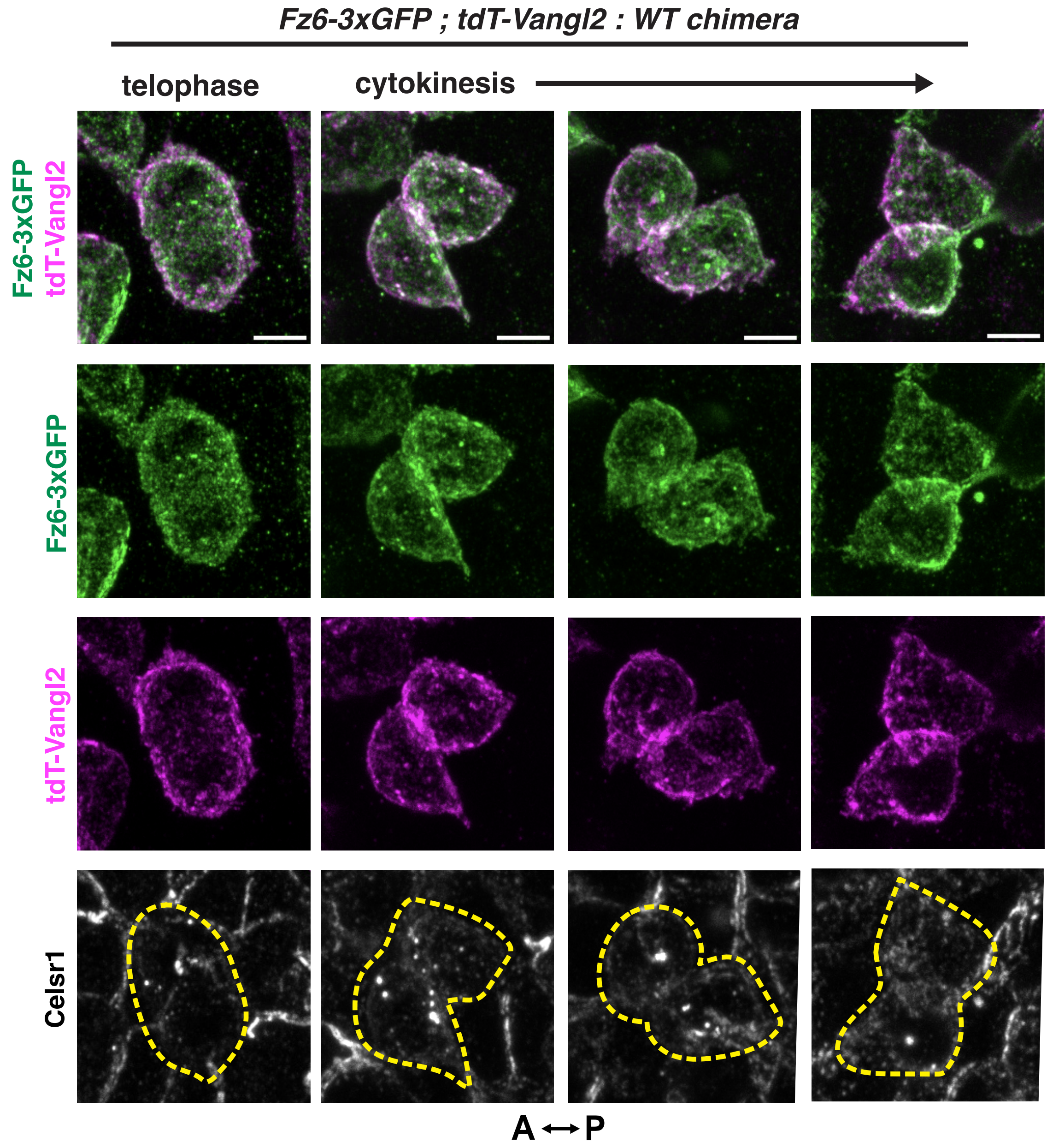


**Supplementary Figure S6. Restoration of PCP protein asymmetry occurs late in cytokinesis.**
Representative examples of isolated basal epidermal cells Fz6-3xGFP (green) and tdT-Vangl2 (magenta) that are completely surrounded by unlabeled wild-type cells in *E*15.5 chimeric mouse embryos (*Fz6-3xGFP;tdT-Vangl2* : *WT*). Chimeric skins are also labeled with Celsr1 antibodies (grayscale). Shown are maximum intensity projections of cells in telophase and cytokinesis (indicated by the presence of Celsr1-containing endosomes). Note the progressive polarization of Vangl2 to the anterior and Fz6 to the posterior of daughter cells. Scale bars, 5 $\mu$m.
